# Postpartum Systemic Inflammation is Reflected in Early Distinct Fecal Microbiome Differences in Dairy Cows

**DOI:** 10.64898/2026.06.12.731891

**Authors:** Lipika Das, Diana G. Puerres Narvaez, Natnicha Taechachokevivat, Alejandro D. Kimball, Rafael C. Neves, Ilya B. Slizovskiy

## Abstract

A tightly regulated inflammatory response occurs during and following parturition; however, excessive or prolonged inflammation negatively affects herd health and productivity. The gut microbiota plays an important role in host immunity and metabolism and undergoes substantial changes during the transition period. However, the relationship between systemic inflammation, measured by serum acute-phase proteins, and gut microbial dynamics during early lactation remains poorly understood. We investigated fecal microbiota dynamics in relation to systemic inflammation in early postpartum dairy cows. Fecal and blood samples were collected from 71 Holstein cows on days 1 and 3 in milk (DIM). The V3–V4 region of the 16S rRNA gene was sequenced and microbial diversity, differential abundance, and microbial interaction networks were evaluated.

Inflammatory status, defined by fibrinogen, haptoglobin, and their combined classification, was associated with significant alterations in fecal microbial composition during the immediate postpartum period, independent of body condition score, parity, DIM, and DNA extraction parameters. Differential abundance analyses revealed extensive taxonomic restructuring, while network analyses identified increased modularity, altered keystone taxa distribution, and greater compartmentalization of microbial interactions in animals with elevated inflammatory markers. Several taxa were consistently associated with inflammatory status across analytical approaches. Notably, *Ruminococcaceae* UCG-002 and *Dielma* were associated with inflammatory states, whereas *Xylanibacter*, *Marvinbryantia*, *Akkermansia*, and *Oscillibacter* were associated with non-inflammatory status.

This study identifies an association between systemic inflammation and fecal microbiota composition in early transition dairy cows, providing a foundation for future microbiome-based biomarkers of inflammatory status.

**IMPORTANCE:** Subclinical systemic inflammation during early postpartum can negatively affect dairy cow health and productivity, yet current monitoring relies on repeated blood sampling and transient inflammatory biomarkers. For the first time association of systemic inflammation, assessed using fibrinogen, haptoglobin, and their combined classification, with alterations in fecal microbial composition, microbial interaction networks, and keystone taxa distribution during the early postpartum period was established. Several bacterial taxa were consistently associated with either elevated or normal inflammatory states across differential abundance, network, and odds ratio analyses. Many of these taxa remain poorly characterized in dairy cattle, highlighting the need for future mechanistic studies. This study demonstrates that systemic inflammation during early postpartum is associated with measurable alterations in the fecal microbiota. This work provides a foundation for developing microbiome-based biomarkers for detecting and monitoring subclinical systemic inflammation in dairy cattle.

## INTRODUCTION

Dairy cows play a fundamental role in global agriculture, providing essential nutrition by generating billions of gallons of milk annually, and supporting a multi-hundred-billion-dollar global industry ((International Dairy Federation, 2025); (U.S. Department of Agriculture, 2025). Health and productivity of herds are significantly affected by events during the periparturient period, one of the most critical stages in a dairy cow’s life marked by physiological and metabolic adjustments necessary for calving and the onset of lactation (Bruinjé & LeBlanc, 2025). During this transition, metabolism is rapidly redirected toward milk synthesis, resulting in markedly elevated energy and protein demands that often exceed voluntary feed intake. Consequently, cows are particularly vulnerable to stressors, frequently leading to inflammatory conditions and metabolic dysfunction (Bradford et al., 2015a). Physiological modifications that parallel these adaptations include reproductive tissue remodeling associated with parturition, mammary gland expansion, uterine tissue trauma in early postpartum, and alterations in gastrointestinal permeability driven by abrupt dietary changes (Sheldon et al., 2009; Baumgard & Rhoads, 2013; Kvidera et al., 2017). Nearly all cows experience negative energy balance due to the high nutritional demands of lactation, along with adipose mobilization, immune activation, and systemic inflammation even in the absence of overt predisposing disease (Serbetci et al., 2024; B.J. Bradford et al., 2015; Mekuriaw, 2023).

While a tightly regulated local and systemic inflammatory response is needed for parturition and postpartum pathogen clearance and tissue repair, extensive inflammatory responses have been associated with impaired productivity and overall health. Even low-grade inflammation measured by higher concentration of acute phase protein has been associated with reductions in milk yield of up to 20%, (Bradford et al., 2015b) as metabolic energy is diverted from lactation and reproduction towards inflammation. Persistent inflammation can also result in serious postpartum complications, including mastitis, metritis, and endometritis, that impair reproductive performance, prolong calving intervals, and increase the risk of premature culling (Bradford et al., 2015b; Brown & Bradford, 2021).

Measuring inflammatory response in dairy cows have been previously reported based on cytokine quantification, including IL-6, IL-10, IFN-γ, and IL-1β (Ishikawa et al., 2004; Trevisi et al., 2015; Westhoff et al., 2025). However, studies by Jahan et al. and Trevisi et al. (2015) have reported only transient and modest cytokine responses around parturition, suggesting that sharply timed inflammatory changes may be missed without high-frequency sampling (Jahan et al., 2025; Trevisi et al., 2015). In contrast, acute phase proteins (APPs) induced by proinflammatory cytokines, including haptoglobin (Hp), fibrinogen (Fb), α1-acid glycoprotein (AGP), inter-α-trypsin inhibitor heavy chain 4 (ITIH4), C-reactive protein (CRP), serum amyloid A (SAA), and lipopolysaccharide-binding protein (LBP) are often preferred in cattle studies because of their longer plasma half-life, which allows for a more stable measure of systemic inflammation (Saco & Bassols, 2023; Reczyńska et al., 2018; Paulina & Stefaniak Tadeusz, 2011). Among these APPs, haptoglobin is considered an especially sensitive and reliable inflammatory biomarker in transition dairy cows (Bionaz et al., 2007; Huzzey et al., 2011). Notably, APPs are expressed at different stages of the inflammatory response and exhibit distinct temporal patterns, underscoring the importance of selecting appropriate markers to accurately capture inflammatory status in various contexts.

Concurrent with metabolic changes in the transition period, the gastrointestinal tract undergoes dramatic structural and functional remodeling. Reduced feed intake around parturition, coupled with increased carbohydrate rich concentrate feeding postpartum, and often insufficient physically effective fiber, frequently leads to ruminal acidosis, a condition characterized by lowered rumen pH that is thought to promote overgrowth of acid-tolerant, Gram-negative bacteria (Khafipour et al., 2009; Saleem et al., 2012). The acidosis-induced altered bacterial landscape has been linked with derangements in ruminal and intestinal epithelium, enteric oxidative stress, tissue catabolism, and ultimately a release of lumenal microbial endotoxins such as lipopolysaccharides (LPS) (Johnzon et al., 2018; Plaizier et al., 2012) which themselves can trigger acute-phase responses and systemic inflammation (Dunière et al., 2021; Guo et al., 2024). Together, emerging evidence supports a gut microbiota–inflammation axis, in which perturbations of the native steady-state microbial communities (i.e. microbiomes) parallel microanatomic alterations of the intestine and systemic immune activation. (Zhao et al., 2024). Most studies to date have focused on overt disease, with less attention given to studying the stability of microbiota in early or subclinical systemic inflammation during the transition period. Moreover, links between gut microbiota composition and inflammation assessed by acute-phase proteins remains largely unexplored, highlighting a critical knowledge gap in identifying microbiota-based biomarkers that may provide more stable indicators of inflammatory status than transient plasma proteins.

Given this knowledge gap, investigating the interactions between systemic inflammation measured using acute-phase proteins and the gut microbiome during the transition period can deepen our knowledge of mechanisms underlying a heightened systemic inflammatory response in the early postpartum. Early identification of inflammatory states could allow timely intervention before inflammation progresses to more severe conditions that negatively impact herd productivity. Our study tests the hypothesis that fecal microbiota composition will vary among animals with elevated or normal markers of systemic inflammation, as defined by blood concentrations of fibrinogen, haptoglobin, and/or their combination. Additionally, we used abundance and network-based correlation strategies to identify specific microbial signatures associated with acute-phase response markers during the early post-parturient period. Ultimately, this study aims to support development of microbiota-based biomarkers enabling early detection of inflammatory sequelae and support improved productivity in dairy cows.

## RESULTS

### Taxonomic profiling was not impacted by sample processing, sequencing, and bioinformatic workflows regardless of systemic inflammation state

A total of 142 fecal samples were collected from 71 nulliparous and multiparous dairy Holstein-cross cows at 1 and 3 DIM (*N*_DIM1_= 71, *N*_DIM3_= 71) between October 2024 and February 2025. Concurrent blood samples were obtained to classify animals into normal or elevated inflammation based on plasma fibrinogen (late-stage marker, cutoff = 0.5g/dL; haptoglobin (early-stage marker; cutoff = 0.45g/L (Kerwin et al., 2022). Inflammation status was assigned only after fecal microbiome collection and sequencing. Haptoglobin levels were measured at 1 and 3 DIM, whereas fibrinogen levels were measured at 1, 3, and 7 DIM. All animals in this study were reared within the same commercial dairy farm (Indiana, USA), and managed under uniform feeding and housing conditions during the sampling period.

To account for potential extra-intestinal contamination, extraction blanks (bead tubes without any fecal sample; *N*= 5), farm-environment control (air sample amended with 1X PBS; *N*= 5); reagent control (1X PBS blank; *N*= 5), and positive controls including MockFecal communities (pooled human microbiota [Zymo Research, Cat No. D6323]; *N*= 3), and MockLog ZymoBIOMICS™ microbial community standard containing 8 bacterial and 2 fungal species [Zymo Research, Cat No. D6311]; (N= 2). Across all libraries, 175,467,458 raw reads were generated targeting the V3-V4 region of the 16S rRNA gene using AVITI sequencing, representing ∼105.3 Gb of total sequenced data. As expected, fecal samples exhibited substantially higher sequencing depth (mean= 586,861; range: 12,821–883,019), and after quality filtering, denoising, merging, and chimera removal with DADA2, a mean 403,813 reads per library remained (range: 6,166–646,836). Mock community libraries similarly yielded high read counts consistent with their known microbial complexity with a mean 785,293 input reads (range: 329,700–995,432) and 628,978 filtered reads retained (range: 286,569–769,729). In contrast, environmental, extraction, and reagent controls received markedly lower sequencing depth and produced fewer filtered, denoised, non-chimeric reads (Figure 1, Supplementary Table S1–S3). Statistical analyses confirmed the strong effect on sample sequencing throughput by sample type (Figure 1C). Fecal samples exhibited microbial community compositions that were strongly and consistently distinct from environmental, extraction, reagent, and mock controls, with large effect sizes (Type III ANOVA pairwise R² = 0.17–0.26, *P*_adj_ < 1e−7). These results are likely supported by parallel findings in the 16S rRNA gene copy numbers found significantly higher in fecal samples, followed by the mock communities (Type III ANOVA (R^2^= 0.16, p < 0.05). In contrast, all non-biological controls exhibited substantially lower copy numbers compared to biological samples (Figure 1B). Read retention rates (i.e., quality filtered, denoised, and chimera removed reads as a fraction of overall raw throughput) were highest for environmental controls (mean: 84.67%) and reagent controls (84.22%), followed by extraction controls (82.47%) and both the mock communities together (81.06%) (Supplementary Table S1). Read retention from fecal samples were more variable, averaging 68.81% (range: 41.63%–76.50%). Following taxonomic assignment of all libraries via the SILVA database (v138.2; Quast et al., 2012), non-target sequences were removed, including 62 chloroplast (order level) and 193 mitochondrial (family level) ASVs classified, resulting in 23,272 ASVs discovered across 162 libraries.

**Figure 1.**
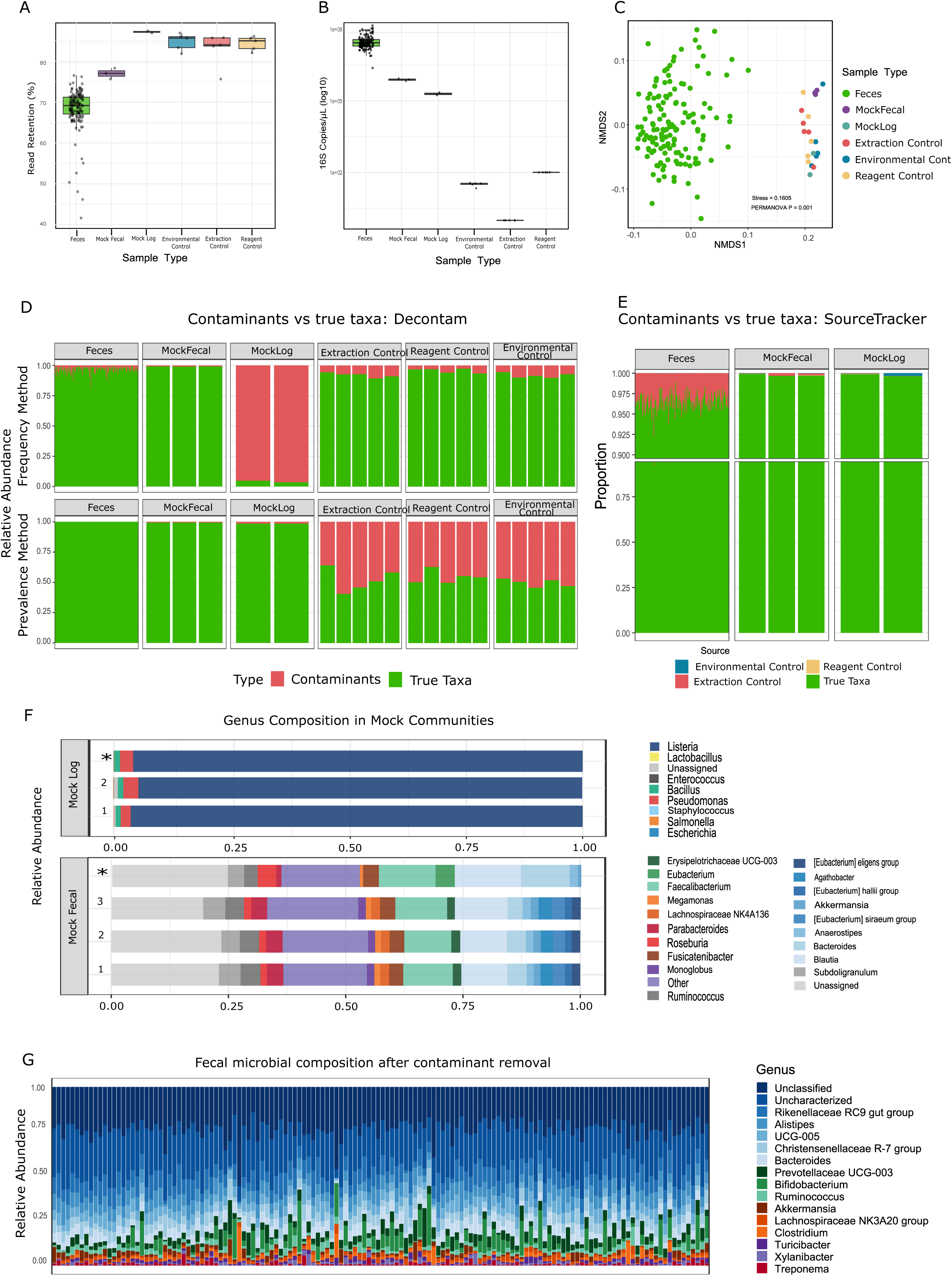
Overview of sample processing, quality control, and microbial composition. **(A)** Read retention rates, representing the percentage of raw reads retained after quality filtering, denoising, and chimera removal across all sample types, including fecal samples, mock communities, and negative controls **(B)** Absolute 16S rRNA gene copies per µL quantified by qPCR, reflecting microbial load differences among sample types. **(C)** Non-metric multidimensional scaling (NMDS) ordination of microbial communities based on Bray-Curtis dissimilarity, illustrating clustering by sample type. **(D, E)** Identification of contaminant ASVs using Decontam (D) and SourceTracker (E), showing contributions from negative controls, reagent controls, and mock communities. **(F)** Genus-level composition of mock community samples (Mock reference (*), MockFecal and MockLog). **(G)** Genus-level composition of fecal samples after removal of contaminant ASVs. Relative abundances are displayed for the most prevalent genera, taxa with less than 1% abundance grouped as Unclassified and taxa lacking genus level assignment are labeled as Uncharacterized.

Despite higher sequencing depth and 16S rRNA gene copy numbers in fecal samples than controls (Figure 1), low-biomass reagent and extraction controls exhibited detectable microbial signals, consistent with technical background contamination commonly reported in amplicon-based microbiome studies (Díaz et al., 2021). To identify and remove potential non-intestinal ASVs introduced during sample handling, DNA extraction, or library preparation, a sequential contaminant-filtering workflow was applied using decontam (Davis et al., 2018) and SourceTracker (Callahan et al., 2016). Putative contaminant sequences were first identified using decontam frequency- and prevalence-based methods, comparing fecal samples with extraction, reagent, and environmental controls. SourceTracker was then used to estimate the contribution of negative controls to fecal microbial profiles. Results from both approaches were integrated to define a final set of contaminant ASVs, which were removed before downstream analysis (Supplementary Figure 2).

Contaminant filtering by decontam R package identified 2,368 unique ASVs as putative non-intestinal contaminants. The frequency method (threshold = 0.5) identified 1,918 ASVs (8.68%), while prevalence-based analyses identified 349 ASVs in reagent controls, 369 in extraction blanks, 335 in environmental controls, and 483 (2.18%) when all negative controls were considered jointly. After merging overlapping detections into a non-redundant contaminant set, these ASVs were removed, reducing the dataset to 20,904 ASVs. Bayesian SourceTracker was then used to estimate contamination from negative controls. Environmental, extraction, and reagent controls were treated as source environments, while fecal samples and mock communities were defined as sinks. Contaminant signals were predominantly derived from extraction controls, accounting for approximately 99% contaminant-associated reads (Figure 1E). Per-sample community composition analyses indicated that most contaminant-associated signals were assigned to *Clostridium, Romboutsia, Turicibacter,* and UCG-005, consistent with the most abundant genus-assigned taxa identified in laboratory extraction kits and dairy farm-borne air (Dean et al., 2024) and laboratory reagents (Agudelo & Miller, 2025). Removal of contaminant ASVs reduced the dataset to 20,810 ASVs across 147 fecal and mock community libraries.

The positive controls generated 5,432,420 raw reads across 3 MockFecal samples (442 MB), and 2,420,514 reads across MockLog samples (146 MB). Comparison with the corresponding Zymo reference databases (https://zymo-files.s3.amazonaws.com/D6323/D6323_NGS_characterization_data.xlsx and https://s3.amazonaws.com/zymo-files/BioPool/ZymoBIOMICS.STD.refseq.v2.zip), identified 1,095 and 45 ASVs in the MockFecal and MockLog communities, respectively. Of the MockFecal ASVs, 953 ASVs, corresponded to the expected community members, whereas the 142 likely represented low level contamination that arose from sample processing or sequencing/analytical noise. Within the MockFecal community, *Bacillota* were the dominant phylum comprising 69.9% of the sequences, though somewhat underrepresented compared to the theoretical 83.3%. *Bacteroidota* were the second most abundant, accounting for 20.6% in our data versus 12.6% theoretical relative abundance, suggesting an overestimation. Other lower-abundance phyla were closely represented in our sequencing relative to expected abundances, including *Actinomycetota* (1.67% vs. 1.44% theoretical), *Pseudomonadota* (1.74% vs. 0.7%), *Verrucomicrobiota* (1.58% vs. 0.73%), *Cyanobacteria* (0.38% vs. 0.29% theoretical), and *Euryarchaeota* (0.02% vs. 0.53% theoretical). Notably, *Lentisphaerota* (0.047%) was present in the theoretical community but not recovered in our sequencing data. This failure to recover may represent a limit of detection following our gDNA extraction and sequencing protocol.

The theoretically expected bacterial composition based on 16S rRNA gene abundance was recovered in the MockLog community, and all expected bacterial taxa were detected. Recovery of expected taxa, *Listeria* (95.6%), *Pseudomonas* (2.71%), *Bacillus* (1.15%), *Escherichia* (0.068%), and *Salmonella* (0.0685%) was consistent with their theoretical abundances (95.9%, 2.8%, 1.2%, 0.069%, and 0.07% respectively). In contrast, *Lactobacillus* (0.00613% observed; 0.012% expected), *Enterococcus* (0.00223%; 0.00067%), and *Staphylococcus* (0.00003%; 0.00010%) showed moderate underrepresentation. Collectively, the mock community results demonstrate that the laboratory workflow reliably recovered expected microbial taxa, supporting the technical integrity of our sequencing methodology. Minor deviations from theoretical relative abundances are consistent with well documented biases arising from limitations in mechanical and enzymatic lysis of Gram-positive bacteria (McLaren et al., 2019) or limitations in sequencing depth of amplicon-based approaches (Jia et al., 2022).

### Compositional differences in fecal microbiota is detected among periparturient dairy cows with early postpartum inflammation

To examine whether fecal microbiota diversity differed between dairy cows with normal and elevated inflammation biomarkers during early postpartum, alpha-diversity was evaluated using four metrics: richness, Shannon diversity, Simpson diversity, and evenness at ASV-level. Linear mixed-effects models were fitted separately for fibrinogen, haptoglobin, and their combined classification (see Methods). Across all models, inflammatory status was not significantly associated with within-sample microbial diversity (all *P*_adj_ ≥ 0.44; Supplementary Table S2), indicating that systemic inflammation was not strongly associated with within-sample diversity. Among covariates, DIM demonstrated the strongest and most consistent association and was negatively associated with all four diversity metrics; however, these did not remain statistically significant after FDR adjustment. Parity, BCS and extraction date were not significantly associated with alpha-diversity (all *P*_adj_ ≥ 0.55) suggesting minimal effects of host characteristics and batch variation on within-sample diversity (Supplementary Datafile 4).

We next assessed whether systemic inflammation was associated with overall differences in fecal microbiota composition using multivariate analyses of beta-diversity based on Bray-Curtis, Aitchison, weighted and unweighted UniFrac distances. Separate models were fitted for fibrinogen, haptoglobin, and their combined classification. Across metrics, inflammatory status explained up to 2.1% of variation in community composition. Fibrinogen status explained 0.7–0.8% of variation (R² = 0.007–0.008) and was associated only with phylogenetic distance based beta-diversity measures (weighted and unweighted UniFrac; *P*_adj_ = 0.0025), with no significant differences detected using abundance-oriented metrics. Haptoglobin status explained 0.6–0.8% of variation and was associated only with abundance-emphasized beta-diversity measures (Bray-Curtis, *P*_adj_ = 0.0025; Aitchison, *P*_adj_ = 0.018), with no significant differences detected using phylogenetic distance metrics. The combined inflammation classification explained the most variation in beta-diversity (1.5–2.1% for Bray-Curtis (*P*_adj_ = 0.004)) and weighted UniFrac (*P*_adj_ = 0.028), which capture both abundance-based and phylogenetic turnover between elevated and normal groups. Beyond inflammatory status, extraction date and DIM consistently explained equal or greater proportions of variation (3.0–7.7% and 0.4–2.0%, respectively; *P*adj ≤ 0.0089), whereas BCS and parity had smaller, metric-dependent effects. Overall, these findings demonstrate that inflammatory status defined by fibrinogen, haptoglobin and their combination is associated with alterations in the fecal microbiota during the early postpartum period (Figure 2). ***Systemic inflammation is associated with broad alterations in abundance of the fecal bacteriome***

**Figure 2.**
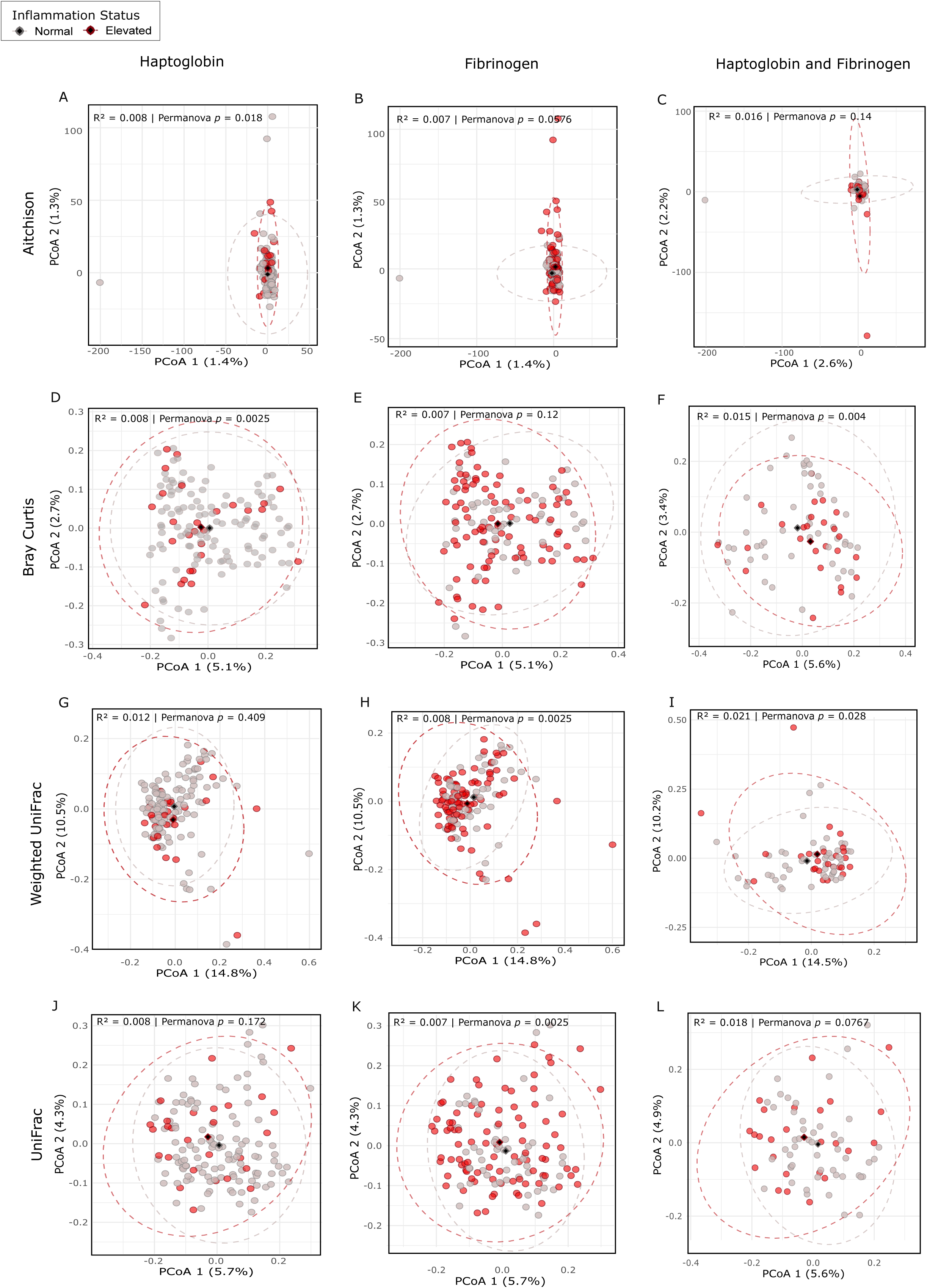
**Beta-diversity in fecal microbiota of dairy cows is distinguishable by inflammation status as defined by different inflammation markers**. Principle coordinates analysis (PCoA) plots are used to visualize differences in fecal microbiota composition among dairy cows identified as having either ‘elevated’ or ‘normal’ inflammation markers, defined by using only serum haptoglobin (**A**, **D**, **G**, **J**), fibrinogen (**B**, **E**, **H**, **K**), or combined haptoglobin-fibrinogen group (**C**, **F**, **I**, **L**). Plots utilize different beta-diversity distance metrics to emphasize different attributes of community diversity, including Aitchison (**A**–**C**), Bray-Curtis (**D**–**F**), weighted UniFrac (**G**–**I**) and unweighted UniFrac (**J**–**L**). Ordinations are summarized by depicting within-group centroids and ellipsoids representing the median and 95% confidence interval, respectively. Significance of ordinated clusters are assessed using a permutational analysis of variance (PERMANOVA) with false discovery rate (FDR) adjustment (*P*_adj_ <0.05), with the proportion of variance explained by each pairwise comparison summarized as *R*^2^.

While beta-diversity analyses identified community-level compositional differences across inflammation states, differential abundance analysis was performed to identify taxa associated with these shifts. Raw ASV counts were modeled in DESeq2 using a negative binomial framework, with inflammatory status as the primary explanatory variable and relevant host and technical covariates included as adjustments (see Methods). Differential abundance was evaluated as log2-fold change (log2FC) using separate models for fibrinogen, haptoglobin, and combined inflammation status. In all models, the normal group served as the reference level, and significance was determined using a Benjamini–Hochberg FDR-adjusted threshold of α = 0.01.

Differential abundance analysis identified extensive microbial shifts across inflammation states. Among 4,834 ASVs shared between elevated and normal haptoglobin groups, 1,452 (30.4%) were differentially abundant, including 891 enriched in the elevated group and 561 differentially abundant in the normal group. Similarly, 1,564 of 5,538 ASVs (28.2%) differed between elevated and normal fibrinogen groups, with 763 enriched in the elevated group and 801 in the normal group. Using a combined inflammation classification, 1,134 of 6,087 ASVs (18.6%) were differentially abundant, including 335 enriched in cows with both markers elevated and 799 enriched in cows with both markers within the normal range. Collectively, these results revealed widespread inflammation-associated shifts in microbial abundance, with numerous taxa exhibiting large log2-fold changes across the inflammation group (Figure 3, Supplementary datafile 6–8).

**Figure 3.**
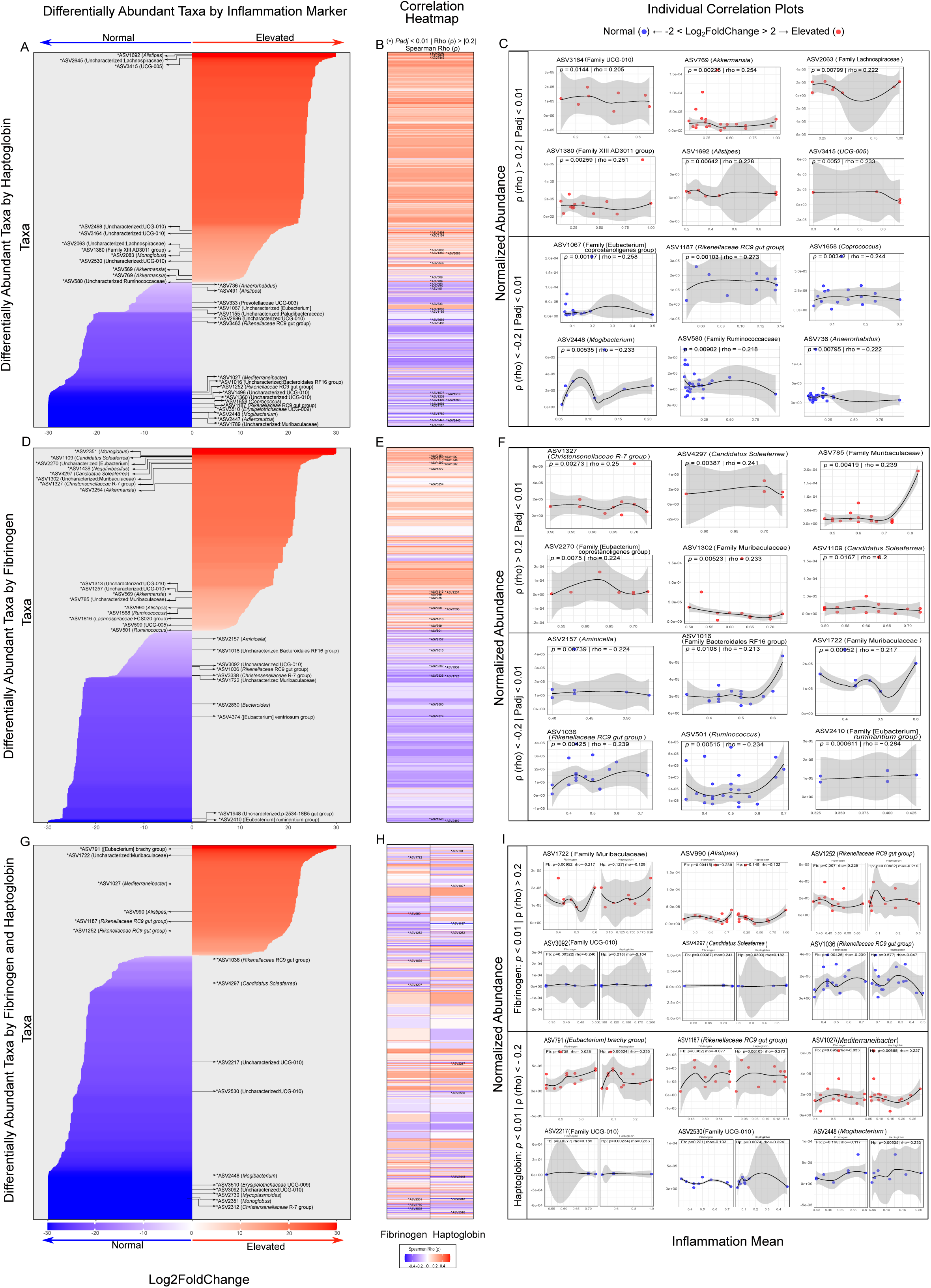
Differential abundance and correlation of fecal microbial ASVs with inflammatory markers. Differentially abundant ASVs associated with serum haptoglobin **(A)**, fibrinogen **(D)**, and combined inflammation status **(G)**. ASVs are shown as horizontal bars centered at zero, with bar length representing log₂ fold change. Positive (red) and negative (blue) values indicate enrichment in elevated and normal inflammation groups, respectively. Highlighted ASVs denote taxa showing significant correlations with the corresponding inflammatory marker (|Spearman ρ| ≥ 0.2). Heatmaps showing Spearman correlations between absolute abundance of differentially abundant ASVs and serum haptoglobin (**B**), fibrinogen (**E**), or combined inflammation status (**H**). Color intensity reflects correlation strength and direction; asterisks (*) indicate *P*_adj_ < 0.01 | Rho (ρ) > |0.2|. Scatter plots illustrate relationships between the serum inflammatory marker concentration and 16S rRNA qPCR-adjusted abundance of the six most positively and six most negatively correlated taxa. **(C, F, I)**. The fitted curve was generated using locally estimated scatterplot smoothing (LOESS; span = 0.9, family = ‘symmetric’), where shared areas represent 95% confidence intervals of fitted mean.

Among genera enriched in cows with elevated serum haptoglobin concentrations were *Alistipes* (log₂FC = 30.0), *Monoglobus* (13.3), and *Akkermansia* (≈ 7.2–10.3), together with members of *Lachnospiraceae* (28.9 and 13.9) and the *Family XIII AD3011 group* (13.5). These taxa are widely recognized as fermentative bacteria capable of degrading complex plant polysaccharides (He et al., 2015). Enrichment of candidate taxa such as UCG-005 (27.2) and UCG-010 (≈ 12.2–17.1) further suggests that systemic inflammation is accompanied by shifts in poorly characterized carbohydrate-utilizing microbiota prominent in the ruminant hindgut ecosystem (Castillo-Lopez et al., 2024). Conversely, several anaerobic gut fermenters and SCFA producing genera were markedly depleted in cows with elevated haptoglobin, including *Erysipelotrichaceae* UCG-009, *Mogibacterium*, and *Adlercreutzia* (log₂FC ≈ −30.0), as well as *Coprococcus* (−29.9), *Mediterraneibacter* (−29.6), and members of the *Rikenellaceae RC9 gut group* (≈ −21.4 to −30.0) (Supplementary datafile 6).

Elevated fibrinogen concentrations were similarly associated with enrichment of several anaerobic fermentative genera, including *Monoglobus* (log₂FC = 26.9), *Candidatus Soleaferrea* (26.6), *Negativibacillus* (24.5), Christensenellaceae R-7 group (22.4), *Akkermansia* (21.9), *Alistipes*, *Ruminococcus*, and Lachnospiraceae FCS020 group (9.7–11.3). These taxa are characteristic members of the mammalian gut microbiota involved in fiber degradation, mucin utilization, and SCFA production. Additional enrichment of UCG-010, UCG-005 (8.5–13.8), *Eubacterium coprostanoligenes* group (25.7), and Muribaculaceae (22.9) suggests links between inflammatory status and microbial carbohydrate, lipid, and host-glycan metabolism. In contrast, cows with elevated fibrinogen exhibited reduced abundance of several fermentative genera, including *Eubacterium ruminantium* group (log₂FC = −30.0), *Eubacterium ventriosum* group (−23.4), *Bacteroides*(−22.9), *Aminicella*, and *Ruminococcus* (−5.6 to −16.7), indicating redistribution within the fiber-fermenting community (Supplementary datafile 7).

When fibrinogen and haptoglobin markers were considered jointly, fewer genera were differentially abundant in cows with concurrent elevations than with those in normal biomarker concentrations. Genera enriched in the elevated group included *Alistipes* (log₂FC = 20.9), *Mediterraneibacter* (22.1), *Eubacterium* brachy group (27.3), and members of the *Muribaculaceae* family (24.8). In contrast, cows with normal inflammatory status displayed enrichment of several fermentative genera including *Monoglobus*, *Christensenellaceae* R-7 group, *Mogibacterium*, and *Candidatus Soleaferrea* (log₂FC ≈ −18.6 to −30.0), together with candidate taxa such as UCG-010 (Supplementary datafile 8).

Overall, elevated fibrinogen and haptoglobin concentrations were associated with taxon-specific shifts in fecal microbiota. To further examine these relationships, correlations between qPCR-adjusted taxon abundance and inflammatory biomarker concentrations were evaluated. Although correlation coefficients varied across taxa (Supplementary datafile 9−11), interpretation focused on taxa showing stronger associations (*P*_adj_ < 0.01; |ρ| ≥ 0.2) (Supplementary Table S3). Haptoglobin concentrations were positively correlated with UCG-010, *Akkermansia*, Family *Lachnospiracaea*, *Family XIII AD3011 group*, *Alistipes*, and UCG-005, and negatively correlated with *Eubacterium* (*coprostanoligenes* group), *Rikenellaceae RC9 gut group*, *Coprococcus*, *Mogibacterium*, Family *Ruminococaceae*, and *Anaerorhabdus*. Fibrinogen concentrations were positively associated with *Christensenellaceae R-7 group*, *Candidatus Soleaferrea*, family *Muribaculaceae*, and *Eubacterium* (*Coprostanoligenes* group). Negative correlations with fibrinogen were identified for *Aminicella*, *Bacteroides RF16 gut group*, *Family Muribaculaceae*, *Rikenellaceae* RC-9 gut group*, Ruminococcus*, and *Eubacterium ruminantium* group. In cows with concurrent elevation of both markers, *Alistipes*, *Candidatus Soleaferrae*, UCG-010 increased with rising inflammation, suggesting their potential utility as microbiota-based indicators of systemic inflammation during early lactation. Conversely, *Rikenellaceae* RC9 gut group, *Mediterraneibacter, Muribaculaceae,* UCG-010, *Eubacterium brachy* group, and *Mogibacterium*, decreased with increasing inflammation, indicating a potentially beneficial role in host immune regulation during the transition period.

### Inflammatory status shapes gut microbial network architecture, keystone module organization, and stability

Microbe–microbe interactions play a critical role in shaping community structure and function thereby influencing host health and productivity. In periparturient dairy cows, systemic inflammation may alter these ecological relationships, highlighting the importance of characterizing microbial network structure across inflammation states and identifying keystone taxa that contribute disproportionately to community stability. To identify these taxa, microbial co-occurrence networks were constructed using the Sparse InversE Covariance Estimation for Ecological Association Inference (SpiecEasi) (v1.1.2) algorithm applied to decontaminated ASV-level phyloseq objects. Networks were inferred separately for each inflammation group: Fibrinogen elevated (FE), Fibrinogen normal (FN), Haptoglobin elevated (HE), Haptoglobin normal (HN), both markers elevated (BE), and both normal (BN). Network topology was accessed using node-level (degree centrality and eigenvector centrality), and network-wide (modularity and assortativity) metrics. Taxa within the top 0.025 percentile for both degree and eigenvector centrality were classified as candidate keystone ASVs, representing highly connected and influential taxa within the microbial community (Supplementary Datafile 12).

Among network associated ASVs, 41 of 3,078 (FE), 14 of 2,230 (FN), 25 of 1,664 (HE), 37 of 3,326 (HN), 17 of 1,604 (BE), and 24 of 2,188 (BN) were classified as keystone taxa. These ASVs spanned 23 taxonomic classes, 14 of which shared across all groups, including *Actinomycetota*, *Bacilli*, *Bacteroidia*, *Clostridia*, *Coriobacteria*, and *Gammaproteobacteria*, representing a conserved core structure. The remaining classes varied among groups, although none were unique to a particular inflammatory state (Supplementary Datafile 12; Supplementary Figure 3). Notably, several highly connected ASVs lacked genus-level annotation, suggesting the presence of uncharacterized yet potentially important community members.

*Bacteroides*, *Rikenellaceae* RC9 *gut group*, *Christensenellaceae* R-7 group, *Lachnospiraceae* (except HE), and UCG-005 (except BE) were recurrent keystone taxa across networks. Keystone taxa detected only in normal networks included UCG-002, *Acetitomaculum*, *Anaeroplasma*, *Anaerorhabdus*, *Atopobium*, *Blautia*, *Dielema*, *Lachnoclostridium*, *Longibaculum*, *Prevotellaceae*, *Ruminococcaceae*, and *Xylanibacter*, with UCG-002 occuring most consistently. Conversely, UCG-010 was a keystone taxon across all elevated inflammation networks. Additional taxa including *Acutalibacter*, CAG-196, Candidatus *Soleaferrea*, Family *Christensenellaceae*, *Escherichia-Shigella*, *Family F082, FD2005, Lachnospiraceae* AC2044 group, *Lachnospiraceae NK4A136 group*, *Monoglobus*, *Olsenella*, Family *Oscillospiraceae*, Family *Peptococcaceae*, Family *Peptostreptococcaceae*, *Prevotellaceae* UCG-004, *Romboutsia*, *Ruminococcus*, *Succinivibrio* were identified as keystone taxa in at least one elevated network, suggesting a role in inflammation-associated community restructuring.

Stronger intra-module connectivity and more compartmentalized microbial interactions were observed under inflammatory conditions. Networks exhibited pronounced modularity (0.47 - 0.77), highest in BE (0.77) and HE (0.75), followed by BN (0.62), FN (0.61), FE (0.53), and HN (0.47). Assortativity showed a similar trend, being highest in BE (0.78) and HE (0.64), intermediate in BN (0.52) and FN (0.50), and lower in FE (0.36) and HN (0.29). Consistent with this organization, keystone ASVs were observed to be non-randomly distributed across network modules and clustered within densely connected regions. Modules enriched with keystone taxa (“*keystone modules*”) likely represent structural units contributing disproportionately to network stability. The number of keystone modules varied across groups (BN: 7, BE: 3, FN: 12, FE: 19, HN: 21, HE: 13; Figure 4), ranging from single-ASV modules to multi-keystone consortia (Supplementary Table S4).

**Figure 4.**
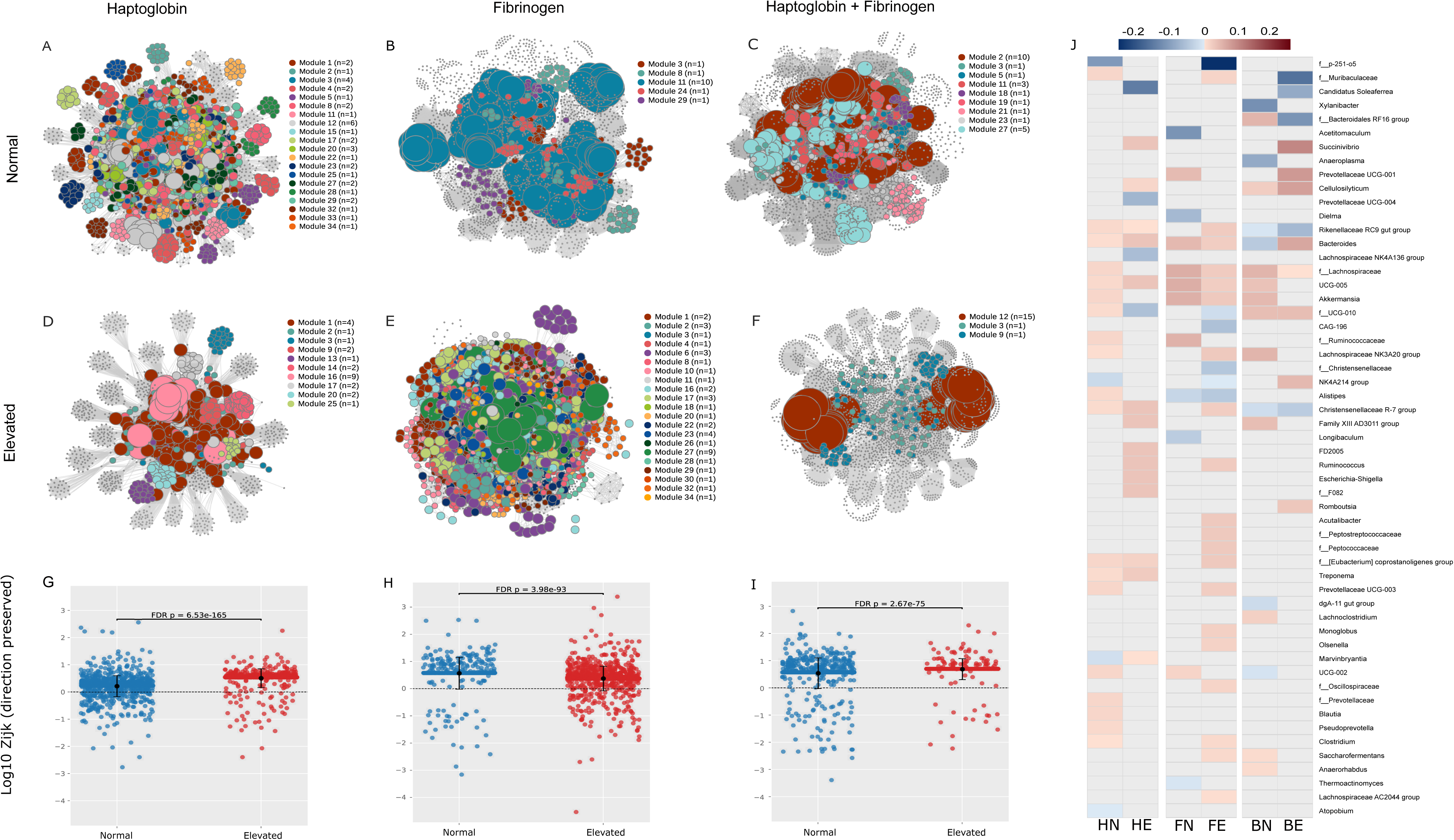
Module-level organization of keystone taxa in microbial co-occurrence networks across inflammatory states. Network visualizations illustrate the distribution and module-level organization of keystone taxa under different inflammatory conditions defined by specific biomarkers, including (**A**) haptoglobin normal, (**B**) fibrinogen normal, (**C**) combined normal, (**D**) haptoglobin elevated, (**E**) fibrinogen elevated, (**F**) combined elevated. Nodes represent microbial taxa defined at the ASV level of taxonomic classification, and edges represent inferred associations between taxa. Modules representing statistically likely network subcommunities containing keystone taxa are highlighted with distinct colors, whereas non-keystone nodes are shown in grey. Node size is proportional to the number of keystone taxa present within each module. The average network interactions of keystone taxa within keystone-driven subcommunities under (**G**) normal vs. elevated haptoglobin, (**H**) normal vs. elevated fibrinogen, and (**I**) normal vs. elevated status for combined group. Statistical significance was assessed using the Mann–Whitney U test with rank-biserial effect sizes and FDR adjustment. (**J**) Summary the estimated propensity for keystone taxa interactions (edges) to occur across subcommunity network modules is presented in a heatmap, where values represent the relative participation of keystone taxa in interactions spanning different modules, aggregated at the genus taxonomic level. Color gradients range from blue to red, indicating negative to positive mean standardized neighborhood participation index (SNPI) values.

To further determine whether keystone taxa organize primarily as network-wide connectors or as module-restricted hubs under different states of inflammation, we evaluated the tendency of keystone bacteria to form co-occurrence interactions with bacteria in keystone modules versus with taxa outside influential modules. These patterns were quantified across the full keystone taxa × module grid within each network, generating a standardized neighborhood participation index (SNPI) enabling direct comparisons of module-level organization. HE and BE networks exhibited significantly higher median SNPI values (*P*_adj_ <0.001), indicating greater clustering of influential taxa and a stronger tendency toward module-restricted organization under elevated markers of systemic inflammation (Figure 5 G–I). This pattern suggests greater possible functional specialization and compartmentalization of network structure. In contrast, animals with elevated serum fibrinogen alone exhibited significantly lower median SNPI, suggesting either a less extensive degree of microbiome re-organization or the preservation of cross-module connectivity characteristic of a more integrated community structure. SNPI analysis revealed group-specific patterns, with keystone bacteria like *Akkermansia*, UCG-005, *Lachnospiracea*, UCG-002 consistently present as module-oriented participants across networks with no systemic inflammation, while *Candidatus Soleaferaee, Ruminococcus,* and *Succinivibrio* were preferentially associated with higher module participation only in animals with elevated inflammatory markers, suggesting possible specialized ecological roles under systemic inflammation and highlighting this latter group of taxa as potential network-based markers of inflammatory states.

**Figure 5:**
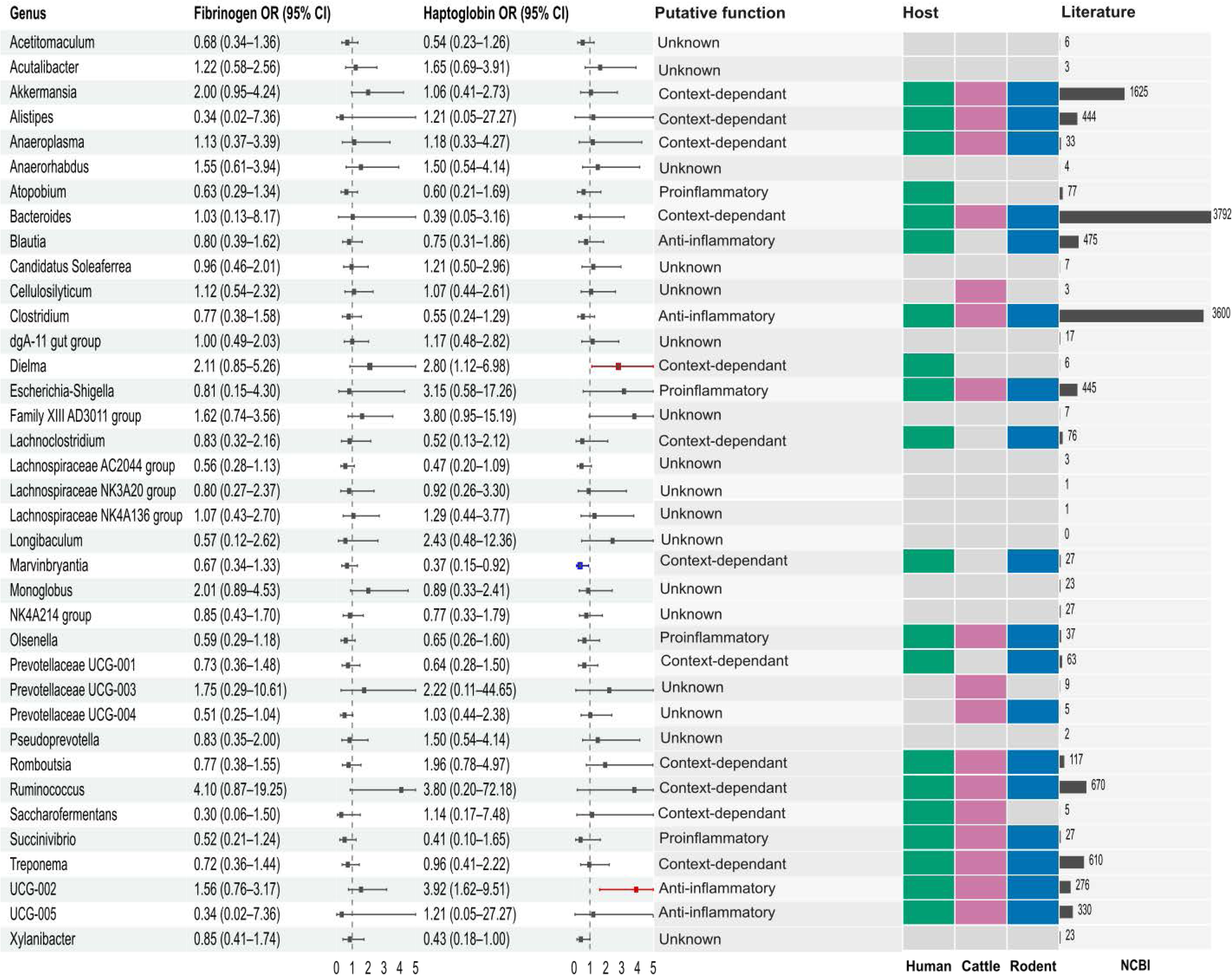
Inflammation associated keystone taxa in fecal microbiome of early postpartum cattle. Forest plots display odds ratios (OR) and 95% confidence intervals (CI) for the presence of keystone genera in relation to elevated inflammation status, defined by fibrinogen and haptoglobin concentrations. Odds ratios were estimated using bias-reduced logistic regression (*brglm2*) with inflammation status as the sole predictor. Points represent OR estimates and horizontal lines denote 95% CIs, with the vertical dashed line indicating no association (OR= 1). Confidence intervals were truncated at the plotting limits for visualization. Keystone taxa are annotated with predicted functional roles (anti-inflammatory, pro-inflammatory, context-dependent, or unknown) based on literature curation. Host-specific evidence (human, cattle, rodent) is indicated by colored tiles, reflecting prior reports linking each taxon to inflammatory processes across different biological systems. The rightmost bar plot represents the number of literature records supporting each taxon–inflammation association, derived from PubMed (NCBI) searches.

To yet further narrow down a candidate list of influential bacteria that may play a critical role in predisposing periparturient animals to systemic inflammation, we quantified the probability likelihoods associated with the presence of any individual network keystone taxon and systemic inflammation using genus-level presence–absence data analyzed with brglm2 (see Methods) (Figure 5; Supplementary Table S5). Several keystone genera exhibited increased occurrence odds in animals with elevated inflammation. Among keystone genera, UCG-002 (OR = 3.92, CI_0.95_: 1.62–9.51) and *Dielma* (OR = 2.80, CI_0.95_: 1.12–6.89) exhibited increased odds under elevated haptoglobin status. In contrast, *Marvinbryantia* exhibited reduced odds under elevated haptoglobin status (OR = 0.37, CI_0.95_: 0.15–0.92). Several additional taxa, including *Ruminococcus* (fibrinogen defined (OR = 4.10, CI_0.95_: 0.87–19.25) and haptoglobin defined (OR = 3.80, CI_0.95_: 0.20–72.18)), displayed elevated odds under inflammatory conditions; however, confidence intervals were wide. In addition to keystone taxa, several non-keystone genera, including *Sphaerochaeta*, Zag_111, *[Eubacterium] siraeum* group, *Butyribacter*, *Anaerovorax*, and *Dorea*, exhibited increased odds of occurrence in inflamed animals despite lacking strong network centrality signatures (Supplementary Table S6).

## DISCUSSION

In the present study, we evaluated association between systemic inflammation and fecal microbiota alterations in Holstein dairy cows during early lactation (DIM 1 and DIM 3) using acute phase proteins Hp and Fb, measured in blood plasma, alongside 16S rRNA gene sequencing. Difference in microbiota compositions were observed between normal and elevated inflammatory states defined by individual biomarkers and their combined classification, indicating that systemic inflammation is associated with measurable shifts in intestinal microbial composition. Notably, both abundance- and phylogeny-influenced beta-diversity metrics showed clearer separation of microbial communities when inflammation status was defined using the combined Hp–Fb marker than individual biomarkers alone (Figure 2), suggesting combined acute-phase marker to be more sensitive to biologically relevant inflammatory states linked to microbiome alterations during the early postpartum period. The transition period is characterized by physiological and metabolic changes to which the intestinal microbiota is highly responsive. Consistent with previous reports, the intestinal microbiota in our study was dominated by *Bacillota*, *Bacteroidota*, *Actinomycetota*, *Pseudomonadota*, and *Verrucomicrobiota* across all animals irrespective of inflammation status. These phyla represent core components of the bovine gastrointestinal microbiome during early life and early lactation (Arnalot et al., 2025; Huang et al., 2020). This leads us to cautiously conclude that most of the inflammation-related alterations to the microbiome occurs through shifts in relative abundance or small-scale phylogenetic changes in specific taxa, rather than large-scale restructuring of core phyla.

Prior studies tracking longitudinal stability in post-partum dairy cow microbiota have reported shifts in fecal microbiota composition over the first few weeks to months following parturition (S. Wang et al., 2025). In contrast, our study demonstrates that significant microbial heterogeneity may be imminently present within the first few days postpartum under states of systemic (i.e. sterile) inflammation, in cows with otherwise clinically inapparent disease. Specifically, we observed significant variation in beta-diversity, differential abundance of ecologically foundational taxa, and network-wide ecosystem differences between groups of cattle as early as Days 1–3 in milk. This early divergence between putatively healthy animals and those determined to have a heightened systemic inflammatory response was not solely attributable to temporal progression, as even after accounting for DIM, sampling time, and extraction date, inflammation status remained a significant determinant of early beta-diversity outcomes (Figure 2, Supplementary datafile 5).

We caution that since blood APPs and fecal microbiota were sampled concurrently, the present study design cannot disentangle the causal and directional relationship between systemic inflammatory status and microbiota composition—that systemic inflammation shapes fecal microbiome remodeling cannot be distinguished from the reverse causal and equally biologically plausible though mechanistically distinct alternative, that pre-existing microbiome states influence host inflammatory responses. Indeed, recent evidence in peri-parturient dairy cattle supports the notion that dysbiosis-induced changes in the gastrointestinal tract can precede the onset of numerous inflammatory and immune-mediated responses of respiratory, reproductive, and mammary systems (Zhong et al., 2026). Similarly, studies in humans have demonstrated bidirectional relationships between inflammation and intestinal microbial composition, whereby chronic inflammatory states can contribute to persistent dysbiosis (Clemente et al., 2018; Ni et al., 2017).

It was noticeable that overall microbial community differences between normal and elevated inflammation biomarker groups were driven by specific taxa and their potential important role and biological relevance in microbial restructuring during systemic inflammation was highlighted across multiple independent analytical approaches, including differential abundance analysis, biomarker correlation, ecological network centrality, module association pattern of keystone taxa, and occurrence-based modeling. Among the most consistently identified genera were *Ruminococcaceae* UCG-002, *Dielma, Ruminococcus, Succinivibrio,* and members of the *Lachnospiraceae* and *Ruminococcaceae* families, collectively characterizing elevated inflammatory states. These taxa were associated not only with compositional shifts, but also with altered ecological influence within microbial interaction networks. In particular, *Ruminococcaceae* UCG-002 and *Dielma* emerged as notable taxa because they were associated with elevated inflammatory networks and additionally exhibited increased occurrence odds under haptoglobin-associated inflammation (**Figure 5**). Previous studies have linked the genus *Ruminococcaceae UCG-002* group to inflammatory and immune-associated disorders, including diffuse large B-cell lymphoma (DLBCL), potentially through modulation of serum monokine induced by gamma interferon (MIG) levels (Jiang et al., 2024). Although mechanistic evidence in cattle remains limited, repeated association of UCG-002 with inflammatory conditions in the present study suggests a potential involvement in inflammation-associated microbial remodeling. Similarly, *Dielma*, a relatively low-abundance anaerobic gut taxon, has previously been associated with bacteremia and chronic kidney disease progression in humans (Lun et al., 2019; Forman-Ankjær et al., 2024). Its enrichment and increased occurrence within inflammatory cattle groups in the present study could reflect adaptation to metabolically altered or inflammatory host environments.

In contrast, taxa associated with normal-inflammatory states, including *Xylanibacter, Marvinbryantia, Akkermansia, Oscillibacter,* and *Salmonella*, appeared to represent microbial features linked to ecological stability and fermentative homeostasis. *Xylanibacter* and *Marvinbryantia* were identified as keystone taxa within animals without elevated inflammation biomarkers, and these keystone bacteria were associated with a reduced likelihood of occurrence in animals with systemic inflammation. These taxa are functionally associated with polysaccharide degradation, fermentative metabolism, and SCFA production, suggesting that maintenance of fiber-fermenting and cross-feeding microbial interactions may contribute to microbiome stability during the postpartum transition period (Kumar et al., 2016; Honda & Littman, 2012; De Filippo et al., 2010). Similarly, *Oscillibacter*, a common gut inhabitant across multiple mammalian hosts including ruminants, is recognized for its involvement in butyrate and valerate production and has additionally been linked to reduction of pro-inflammatory lipid metabolites and improved host metabolic health (C. Li et al., 2024). *Akkermansia*, although exhibiting context-dependent patterns across analyses, has been widely reported to possess anti-inflammatory and gut barrier-supportive properties and has been explored as a probiotic candidate in several inflammatory disease models (Li et al., 2019; Zhang et al., 2025). Interestingly, the increased occurrence of *Salmonella* within normal-inflammatory groups in the present study raises important questions regarding its ecological role within the cattle gastrointestinal microbiome. *Salmonella* is classically recognized as an enteric pathogen associated with diarrhea, septicemia, and respiratory disease in cattle; however, asymptomatic carriage and long-term intestinal persistence are also common in clinically healthy ruminants (Gragg et al., 2013; Stipetic et al., 2016; Kagambèga et al., 2013).

Importantly, several non-keystone taxa, *Sphaerochaeta*, Zag_111, [*Eubacterium*] siraeum group, *Butyribacter*, *Anaerovorax*, and *Dorea*, demonstrated elevated odds of occurrence in inflamed animals despite lacking strong network centrality signatures (Supplementary Table S6). These findings suggest that inflammation-associated microbiome alterations extend beyond highly connected keystone taxa and may also involve opportunistic or inflammation-responsive community members. Although the functional roles of these genera within the postpartum bovine gut remain incompletely understood, previous studies have linked them to inflammatory conditions, mucosal remodeling, host metabolic regulation, and SCFA production (Murray et al., 2002; Wang et al., 2023; Ku et al., 2025; C. Zhang et al., 2023; Liu et al., 2024).

Despite the stated findings of this study, several limitations should be considered when interpreting the results. First, approximately 20% of taxa could not be assigned at the genus level and species level indicating a limited taxonomic resolution of 16S rRNA sequencing. Notably, several differentially abundant ASVs and keystone bacteria remained taxonomically unresolved, reflecting limitations in current reference databases and annotation frameworks for 16S rRNA amplicons. In addition, the use of 16S rRNA gene sequencing inherently does not permit direct functional inference of specific genetic content. Nonetheless, 16S-based approaches remain a cost-effective and widely adopted standard for large-scale microbiome studies, enabling robust comparative analyses of community structure across biological conditions. Second, the use of fecal microbiota as a proxy for intestinal and ruminal microbial communities. While this approach does not fully capture compartment-specific bacterial dynamics, the observation that fecal microbial composition varies with systemic inflammatory status suggests that our findings may represent a conservative estimate of microbiome perturbations occurring along the gastrointestinal tract. Third, sampling was restricted to two early postpartum time points (DIM 1 and 3), limiting temporal dynamics of systemic inflammation and microbiome transitions. Although higher-frequency sampling could provide finer resolution of early, mid, and late inflammatory phases, such approaches are often impractical and ethically challenging in herd-based studies. Importantly, our sampling framework aligns with established biomarker monitoring practices during the transition period, ensuring relevance to real-world dairy management systems. Finally, systemic inflammatory markers were not directly linked to gastrointestinal-specific inflammation or circulating microbial signatures. While serum haptoglobin and fibrinogen were used to classify inflammatory status, markers of intestinal inflammation (e.g., fecal calprotectin or lactoferrin) and blood-borne microbiota were not assessed. These intestinal biomarkers and their assay systems have not been validated for use in dairy cows, though at the time of manuscript preparation, a recent report points to potential opportunities for their use (Saco et al., 2025). Incorporating these measures in future studies would strengthen causal inference and help validate mechanisms such as increased intestinal permeability (i.e. “leaky gut” model) contributing to systemic inflammation. Ongoing efforts to validate these intestinal biomarkers will further enhance interpretation of host–microbiome interactions.

## CONCLUSION

This study demonstrates that systemic inflammation in the periparturient period, as assessed using fibrinogen and haptoglobin, is associated with alterations in the fecal microbiota of early postpartum dairy cows. Microbial community composition differed between inflamed and normal animals, with distinct patterns captured across diversity and abundance metrics. Several taxa were differentially abundant between groups, and some were significantly correlated with inflammation markers, suggesting their potential as early microbial indicators of systemic inflammation. Network analyses further revealed reorganization of microbial interactions under inflammatory conditions, characterized by redistribution of keystone taxa into fewer modules and increased clustering of functionally similar taxa. Their non-random distribution and differential participation across inflammation states, supported by standardized neighborhood participation index patterns, indicate a structured, inflammation-associated reconfiguration of the microbial network. Interestingly, while several taxa identified in this study have been previously associated with inflammation across cattle and other host systems (including humans and rodents), many lacked prior evidence of such associations, highlighting their potential as novel inflammation-associated microbial signatures.

Collectively, this study identifies the early postpartum period as a critical window of host–microbiome interaction and provides a foundation for developing microbiome-based biomarkers for the early detection of subclinical systemic inflammation in dairy cattle.

## MATERIALS AND METHODS

### Animal Population Selection and Sample Collection

All experimental procedures involving animal use were reviewed and approved by the Purdue University Animal Care and Use Committee (Protocol # 0123002347). The study was conducted at a single commercial dairy farm located in Northwestern Indiana, USA. Between October 2024 and February 2025, a total of 71 Holstein dairy cows were enrolled in this study. While nulliparous cows were kept in a separate dry lot pen from multiparous cows before expected calving and commingled in a different dry lot pen at pre-parturition, all cows were then moved to a single free-stall pen just prior to the sampling phase. All animals were fed *ad libitum* total mixed rations.

Each animal was sampled twice, once on day 1 in milk (DIM1) and again on day 3 in milk (DIM3) for feces, and thrice each on DIM1, DIM3 and DIM7 for blood. All animals were apparently healthy on veterinary examination at the time of enrollment and sampling, and were not administered any chemotherapeutics during the sampling phase. On retrospective review of herd health records, of the 71 enrolled animals, one cow was recorded to have ketosis at day 6 in milk and two animals had metritis after day 5 in milk. Given the low prevalence of these later herd health events, no animals/samples were excluded from intended analysis.

### Sample Collection

Fecal samples were aseptically collected directly from the rectum using sterile obstetric sleeves to prevent environmental contamination. Blood samples were obtained via coccygeal (tail vein) venipuncture using a 20-gauge × 2.54 cm needle into EDTA-containing and lithium heparin-containing vacuum tubes following morning milking to assess systemic inflammatory status using plasma fibrinogen and serum haptoglobin concentrations. An additional air-purged, empty vacutainer was included as an environmental control (n = 5). Following collection, all samples were immediately stored at 4°C and transported to the laboratory on ice. Fecal samples were stored at −80 °C until further analysis. Blood samples were centrifuged at 2,000 × *g* for 10 min at 4 °C. EDTA plasma was analyzed for Fb using a modified heat precipitation method (Millar et al., 1971). Initial plasma protein concentration was measured using a digital refractometer (Misco, Solon, OH). Then, 500 μL of plasma was transferred to a microcentrifuge tube, heated at 56 °C for 5 min to precipitate Fb, and centrifuged at 8,000 × *g* for 10 min. Plasma protein concentration of the supernatant was measured using the digital refractometer. Fibrinogen concentration was estimated by subtracting post-precipitation plasma protein concentration from the initial plasma protein concentration. Heparinized plasma samples were stored at −80 °C until further analysis. Variables including animal identification (Cow ID), body condition score (BCS), and parity were recorded for each cow at the time of sampling. Systemic inflammation was evaluated using plasma fibrinogen and serum haptoglobin concentrations. Plasma fibrinogen was determined using the heat precipitation method, while plasma haptoglobin was quantified using a bovine haptoglobin ELISA kit (Life Diagnostics, West Chester, PA) according to the manufacturer’s instructions. Heparinized plasma samples at DIM1 and DIM3 were diluted 5,000-fold and analyzed in duplicates. Standards were analyzed in triplicates. Intra- and inter-assay coefficients of variation were 2.8% and 11.8%, respectively. These variables were recorded for each individual and used for downstream data analysis and metadata integration (Supplementary Table 1).

### DNA Extraction and Sample Preparation

Total genomic DNA was extracted from all fecal samples using the PowerSoil Pro Kit (Qiagen, Cat. No. 47016, Hilden, Germany), following the manufacturer’s protocol with slight modifications. Briefly, 50 mL centrifuge tubes containing fecal samples were surface-sterilized by spraying with 70% ethanol and allowed to thaw inside a laminar flow hood for approximately 1 hour. A total of 250 mg of fecal material was weighed under sterile conditions and transferred into PowerBead Pro tubes containing 800 μL of Solution CD1. Mechanical lysis was performed using the MP FastPrep-24™ 5G Tissue Homogenizer (MP Biomedicals) under the instrument’s default settings for soil samples. Environmental control tubes, consisting of air-purged, empty vacutainers, were washed with 1 mL of sterile phosphate-buffered saline (PBS), and the recovered PBS was processed for DNA extraction alongside the biological samples using the same protocol.

DNA extraction was conducted in batches of 23-29 samples along with positive and negative control. The extraction blank control (n = 5) consisted of 800 μL of Solution CD1 in an empty PowerBead Pro tube without any sample material, used to monitor for potential contamination from the DNA extraction kit (termed as extraction control in the study). Two positive control communities were included to assess extraction efficiency and taxonomic accuracy: Mock Fecal, the Fecal Reference with TruMatrix™ Technology (Zymo Research, Cat. No. D6323), and MockLog, a defined microbial community composed of eight bacterial and two fungal species (ZymoBIOMICS™ Microbial Community Standard, Zymo Research, Cat. No. D6311). DNA from all fecal samples, MockFecal and MockLog, environmental controls, and extraction blank controls was eluted in 60 μL of TE buffer and the concentration and purity of extracted DNA were assessed using both Nanodrop UV/VIS spectrophotometry (Thermo Fisher Scientific, USA) and fluorometric quantification via PicoGreen staining (BioTek Instruments, USA).

### Quantification of 16S rRNA Gene Copy Number by qPCR

To quantify the total bacterial load, 16S rRNA gene copy numbers were determined using quantitative PCR (qPCR) at UMGC facility. DNA extracted from fecal samples was used as the template to amplify the V4 region of the 16S rRNA gene using the identical primers listed above. Template DNA was used to create 1:8 and 1:64 dilutions from each sample, and all samples were loaded and run on 384-well plates. Previously generated standard curves were used to calculate the amount of material present in each sample. qPCR was run using 6 ul volumes; each reaction contained 3 ul of template and 3 ul of KAPA HiFi mastermix on a Qiagen QuantStudio 5 thermocycler. Amplification conditions consisted of 5 min at 95 °C, followed by 35 cycles of 20s at 98 °C, 15 s at 55 °C, and then 1:00 m at 72 °C.

### Microbiome sequencing

To characterize the bacterial sample composition, the V3-V4 region of the bacterial 16S rRNA gene was amplified using the primer pair V4_515F_Nextera (TCGTCGGCAGCGTCAGATGTGTATAAGAGACAGGTGCCAGCMGCCGCGGTAA) and V4_806R_Nextera (GTCTCGTGGGCTCGGAGATGTGTATAAGAGACAGGGACTACHVGGGTWTCTAAT) for library preparation. Samples were all uniquely dual indexed as detailed in (Gohl et al., 2016). Sample abundances were normalized using SequalPrep kits (Invitrogen), pooled into libraries, and cleaned with AMPure XP mag beads (Beckman Coulter). Final amplicon size was confirmed by testing an aliquot of the final library with an Agilent D1000 TapeStation instrument (Agilent). Prepared amplicon libraries were sequenced on an Element Biosciences AVITI™ sequencer using a 2x300 Medium Run reagent kit, targeting an expected sequencing depth of approximately 588,000 paired end reads per sample.

### Bioinformatics and Sequence analysis

All bioinformatic analyses were performed in R statistical software (v4.4.3; R Core Team 2025; https://www.r-project.org/), and visualizations were generated using the ggplot2 package (v3.4.2) (Wickham, 2016). For quality control paired-end sequence reads were evaluated for quality and adapter contamination using FastQC (v0.11.7) (Andrews, 2010). Reads were then processed with Trimmomatic (v0.39) (Bolger et al., 2014) to remove low-quality bases and sequencing adapters. Trimming parameters included removal of Nextera adapter sequences, cropping of the first 20 bases from the 5′ end, trimming low-quality bases from both ends, applying a sliding window filter, and discarding reads shorter than 50 bp. Only properly paired reads that passed all quality control steps were retained for downstream analysis using DADA2 (Callahan et al., 2016). The resulting trimmed reads were further processed using the DADA2 pipeline (v1.34.0) for quality filtering, error correction, and microbial community inference. The filterAndTrim function in DADA2 was used to remove low-quality reads and prepare the data for denoising. Forward and reverse reads were truncated to 240 bp and 220 bp, respectively, based on their quality score profiles. To remove residual primers, the first 20 bases of the forward reads and 17 bases of the reverse reads were trimmed using the trimLeft parameter. Reads containing ambiguous base calls or matching PhiX sequences were discarded. Additionally, only reads with a maximum expected error rate of ≤3 for forward reads and ≤4 for reverse reads were retained for downstream processing.

The learnErrors function was used to estimate error rates specific to the Element Biosciences AVITI™ sequencer, which were subsequently used to correct sequencing errors. Reads that were either shorter or longer than the expected amplicon length were removed. Forward and reverse reads were then merged using the mergePairs function, and chimeric sequences were identified and removed using removeBimeraDenovo. Merged reads longer than 250 bp or 258 bp were discarded based on the distribution of read lengths and the expected size of the amplified V4 region. Following the removal of chimeric sequences, the resulting amplicon sequence variants (ASVs) were partitioned based on sample type to utilize the most appropriate reference databases for taxonomic assignment, which was executed using the *assignTaxonomy* function. Fecal and technical control ASVs were aligned and classified against the SILVA reference database (v138.2) (Quast et al., 2012). Conversely, the MockFecal and MockLog ASVs were segregated and classified only against their respective, specific mock community reference databases ((https://zymo-files.s3.amazonaws.com/D6323/D6323_NGS_characterization_data.xlsx) and (https://s3.amazonaws.com/zymo-files/BioPool/ZymoBIOMICS.STD.refseq.v2.zip)) to provide an accurate measure of technical classification efficacy. Additionally, non-bacterial sequences classified as chloroplast or mitochondrial in origin were removed from the taxonomy table to ensure that only relevant bacterial ASVs were retained for microbiome analysis. The ASV abundance matrix, taxonomy table and sample metadata were used to generate a phyloseq object for microbiome data analysis using the phyloseq (v 1.50.0) (McMurdie & Holmes, 2013) R package (v 4.4.3). To integrate sequencing statistics with sample metadata, the read tracking summary was merged with the original metadata file, allowing downstream analysis to incorporate read retention information at each step of processing.

### Removal of potential contamination

To identify and remove potential contaminants introduced during sampling, DNA extraction, and sequencing, we employed both the decontam (v1.14.0) (Davis et al., 2018) and SourceTracker (v1.0) (Knights et al., 2011) packages. The initial screening for contaminants was performed using the decontam package on the phyloseq object. Contaminant detection was carried out using both frequency-based and prevalence-based methods via the *isContaminant*() function, with a classification threshold of 0.5, following recommendations by (Davis et al., 2018). For the frequency-based method, potential contaminant ASVs were identified by plotting their abundance against 16S rRNA gene copy number (quantified via qPCR) using the *plot_frequency()* function. ASVs classified as contaminants by this method were removed prior to the prevalence-based analysis. The prevalence-based method was conducted in three iterations, each using a different set of negative controls to capture contaminants specific to distinct stages of sample processing:

1. extraction controls, (2) reagent controls, and (3) environmental controls. The results from all three runs were combined to create a final decontaminated phyloseq object, which was then used for subsequent SourceTracker analysis.

To further estimate and eliminate contamination from the fecal samples, we applied the SourceTracker(v1.0) package. Positive control samples were treated separately for this analysis. Extraction, reagent, and environmental controls were designated as source environments, while the fecal samples were treated as sink environments. SourceTracker predicted the proportional contribution of each source to the ASVs present in each fecal sample. ASVs attributed to contamination were removed, allowing recovery of the true microbial community structure. This dual-step approach produced a refined, decontaminated phyloseq object, which was subsequently used for all downstream microbial community analyses.

### Mock Community Sequencing Analysis

Raw 16S rRNA sequences from two different mock communities, including MockLog-ZymoBIOMICS Microbial Community DNA Standard II (Log Distribution), which contains genomic DNA from eight bacterial and two fungal strains, as well as MockFecal- ZymoBIOMICS Fecal Reference with standard fecal community standard, were analyzed alongside DNA extraction negative controls. To assess contamination introduced during DNA extraction and sequencing, the taxonomic composition of the mock communities (positive controls) was compared against their known reference compositions. Amplicon sequence variants (ASVs) present in the mock communities that did not match the expected reference sequences and were also found in the negative controls at a relative abundance greater than 10% were classified as contaminants and removed from the dataset.

The sequence data for the positive control sample were aligned to the manufacturer’s sequence database (https://s3.amazonaws.com/zymo-files/BioPool/ZymoBIOMICS.STD.refseq.v2.zip) containing 16S rRNA reference sequences for each mock bacterium. For the ZymoBIOMICS Fecal Reference, raw 16S rRNA sequencing reads were obtained from the Zymo Research website (https://zymo-files.s3.amazonaws.com/D6323/D6323_NGS_characterization_data.xlsx). To account for differences in microbial load in different samples, the raw ASVs were normalized with absolute 16s qPCR copynumber. We used a generalized linear model (GLM) to assess differences in 16S rRNA gene copy numbers among sample types, with log-transformed copy number as the response variable and sample type as the predictor. Samples were categorized into three inflammation groups based on inflammatory biomarkers: fibrinogen normal and elevated (cutoff- 0.4g/dL), haptoglobin normal and elevated (cutoff- 0.45g/L), and a combined fibrinogen–haptoglobin classification. In the combined group, samples were defined as normal when both biomarkers were within their respective reference ranges and as elevated when both fibrinogen and haptoglobin concentrations exceeded their defined cutoff values. All the microbial composition and interaction analysis were performed for each of these inflammation groups.

### Microbiome diversity analysis

Four alpha-diversity metrics were employed to evaluate within-sample microbial diversity: Observed richness (number of amplicon sequence variants, ASVs), Shannon diversity index (incorporating richness and evenness), Simpson index (accounting for dominance), and Pielou’s evenness. Observed, Shannon, and Simpson indices were computed using the *estimate_richness()* function from the *phyloseq* R package (v1.50..0), while evenness was calculated using the *evenness()* function from the *microbiome* package (v1.22.0) (Leo Lahti [Aut, 2017). Associations between alpha-diversity and covariates were evaluated using mixed linear regression models implemented via the base lmer::Test() function in R. The models included continuous predictors: fibrinogen and haptoglobin inflammation status, parity grouped into <=3 and >=3, body condition score (BCS) grouped into <=2.5, 3.5 and >=3.5, and days in milk (DIM), extraction date. Only samples with complete metadata for all covariates were included in the analysis. Model fit was assessed using the coefficient of determination (R²), and *P*-values for individual predictors were adjusted using the Benjamini–Hochberg false discovery rate (FDR) procedure. Predictors with adjusted *P*-values (i.e., q-values) below 0.05 were considered statistically significant.

Beta-diversity was assessed using principal coordinates analysis (PCoA) on multiple distance matrices. For community composition, Bray-Curtis dissimilarity and Aitchison distance were calculated; for phylogenetic compositional differences, weighted and unweighted UniFrac distances were computed. Group differences were tested using PERMANOVA (*adonis2*, vegan package) (Oksanen et al., 2001) with 999 permutations. The same covariates included in the alpha-diversity models were considered here.

### Quantification of differentially abundant taxa and taxon–biomarker correlation analysis

In early postpartum cattle, the fecal microbiota alternation is not only influenced by the inflammation measured by inflammation marker itself but also by host health status (BCS), parity (primiparous or multiparous) even individual variation also contribute to shaping microbial abundances within groups. To identify taxa that were differentially abundant between inflammation status groups, we used DeSeq2 (v1.46.0) (Love et al., 2014) to determine the differentially abundant taxa in all the cattle measured by both fibrinogen- and haptoglobin-based markers.

To identify differentially abundant taxa associated with inflammation status, we performed count-based modeling using the negative binomial distribution (NBD)–based package DESeq2 . The count matrix was extracted from the phyloseq object, and taxa with zero counts across all samples were removed prior to analysis. The resulting count matrix (ASV-level) was combined with the sample metadata into a DESeq2 dataset object, with inflammation status (measured by blood haptoglobin or fibrinogen concentration; Normal vs. Elevated) as the primary variable of interest. BCS, parity, and batch extraction date were included in the design formula as covariates to account for potential confounding effects. Counts normalization was carried out using the *estimateSizeFactors()* function and the poscounts method, which is well suited for sparse microbial count data. Differential abundance testing was performed using the Wald test within the DESeq2 framework. Statistical significance (q-value) was set as an FDR-adjusted *P*-value via Benjamini–Hochberg correction below 0.01. Taxa meeting this threshold were considered significantly differentially abundant between inflammation states. For interpretation, both the log_2_-fold change (‘log_2_FC ’) and the corresponding adjusted *P*-value were considered to identify the most strongly differentially abundant taxa. Potential outliers were evaluated using Cook’s distance as implemented in DESeq2. Samples exhibiting unusually large deviations in taxon counts relative to model expectations were flagged as influential observations, and taxa containing such extreme count outliers had their associated *P*-values and adjusted *P*-values set to NA to minimize the risk of spurious detection of differential abundance.

Correlation of differential abundant taxa with inflammation marker concentration was defined by spearman correlation. The taxa abundances were normalized by their corresponding 16S rRNA gene copy numbers, and the inflammation concentration was defined as the mean of the inflammation score measured at DIM1 and DIM3 for haptoglobin and DIM1, DIM3 and DIM7 for fibrinogen. *P*-values were adjusted using the Benjamini–Hochberg (BH) procedure to control the False Discovery Rate (FDR) at a q<0.01 threshold. Taxa with a rho value more than **±** 0.2 and a and a q-value below 0.01 were considered for exploration. Taxa above the cutoff were plotted. The fitted curve was generated using locally estimated scatterplot smoothing (LOESS; span = 0.9, family = ‘symmetric’), where shared areas represent 95% confidence intervals of fitted mean.

### Microbial co-occurrence network analysis

Microbial communities are complex and highly interconnected, and this complex interaction defined by taxonomic co-occurrence has been linked with significant host health outcomes (Kodera et al., 2022). Thus, to detect alterations in co-occurrence patterns associated with systemic inflammation status, microbial networks were inferred separately for each serum inflammation assay using the SparCC-based sparse inverse covariance estimation implemented in the SpiecEasi package (v1.1.2) in R (Kurtz et al., 2015). Network inference was performed on decontaminated ASV-level phyloseq objects for the following groups: both elevated (BE), both normal (BN), fibrinogen elevated (FE), fibrinogen normal (FN), haptoglobin elevated (HE), and haptoglobin normal (HN). Taxa with zero total counts across samples were removed prior to analysis. Networks were inferred using the Meinshausen–Bühlmann (MB) neighborhood selection method to estimate sparse inverse covariance matrices, accounting for the compositional nature of microbiome data. A regularization path of 20 lambda values was used, and model selection was performed using the Stability Approach to Regularization Selection (StARS) with 50 subsampling repetitions. Final co-occurrence networks were constructed using an undirected, unweighted graph *G*_n_ = (*V*_n_, *E*_n_) for a given network *n*, where *V*_n_is the set of ASVs and *E*_n_is the set of edges representing significant covariance associations.

Network topology was evaluated using both node-level and network-level metrics. For node-level metrics, degree centrality and eigenvector centrality were calculated to assess the local connectivity and global influence of individual taxa within the network. Network connectivity was assessed using the igraph package and all inferred networks consisted of a single connected component with no isolated nodes; therefore, no subgraphs or disconnected components required separate handling prior to centrality analyses. For network-level properties, modularity and assortativity were computed to characterize overall network organization. Modularity was calculated based on Louvain community detection. Nominal assortativity was computed using module membership as the node attribute to quantify the tendency of taxa to preferentially interact within modules. Assortativity was calculated based on the module mixing matrix following the formulation of (Newman, 2003) as implemented in the assortativity_nominal() function in the igraph (v2.2.2) package.

Specifically for modularity analysis, each ASV taxon node 𝜏 ∈ *V*_n_was assigned to a putative subcommunity ‘module’ using the Louvain community detection algorithm (Blondel et al., 2008), such that a set of *m*(𝜏) ∈ {1, . . ., *M*_n_} is constructed for each *G*_n_. Additionally, ASVs whose covariance with other taxa predicts putative importance for overall network architecture, defined as ‘keystone taxa’, was profiled by identifying ASVs ranking within the top 2.5th percentile for both degree(𝜏) and eigenvector centrality (𝜏) for each network, establishing a set of *KT*_n_ ⊆ *V*_n_. As for all other modular networks, keystone taxa are presumed to be connected based on the ecological role fulfilled in a complex set of ecological interactions (Guimerà & Nunes Amaral, 2005). We hypothesized that alterations to the extent keystone taxa participate within a given set of modules or sub-compartments of networks indicates a remodeling of their ecological role, and that a global increase in keystone-module interactions compared to the overall distribution of keystone interactions will differ in ecological networks derived from animals with normal vs. elevated serum inflammation biomarkers. To test this hypothesis, for each keystone ASV 𝜏 ∈ *KT*_n_, we quantified the number of edges from each keystone 𝜏 to nodes belonging to module *k* defined as 𝜆_rk_, and the total keystone 𝜏 total number of edges of *G*_n_ defined as 𝜆_r_ using an indicator variable for each, calculated explicitly as:

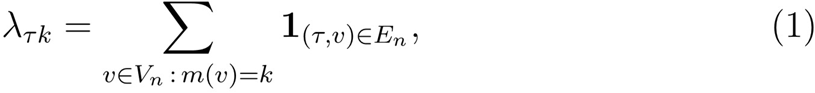

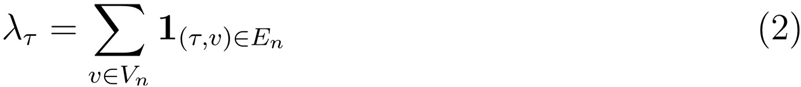

With these quantities, we constructed a normalized participation score *P*_rnk_representing the fraction of each keystone taxon’s total network connectivity directed within modules of a given network:

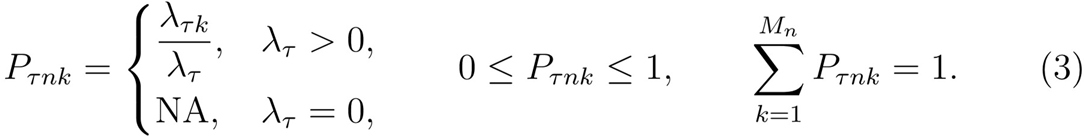

To enable comparison of participation patterns across keystone taxa within the same network, we standardized neighborhood participation values across the full keystone taxa x module grid space, which we refer to as the *standardized neighborhood participation index* (SNPI).We define the SNPI function (*Z*_rnk_) based on the global network-specific 𝜇_n_ and 𝜎_n_:

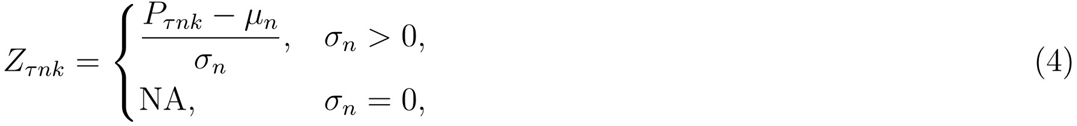

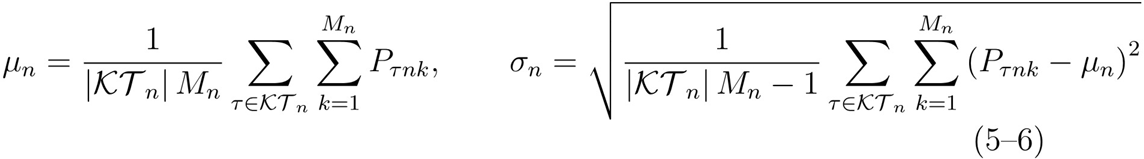

Positive *Z*_rnk_values indicate disproportionately strong targeting of a given module by a keystone taxon relative to the network background. Analysis of altered keystone regionalization patterns between inflammation states were performed by analyzing differences in the mean non-zero log_10_-transformed SNPIs using the Mann–Whitney U test. Significance was assessed at *P*_adj_ < 0.05 and ranked effect size >0.3 or < −0.3. All network metrics were calculated using the *igraph* package (v 2.0.0) in R (Antonov et al., 2023).

### Association and Functional Characterization of Keystone Taxa

To further evaluate the association between keystone taxa and systemic inflammation status, the phyloseq object was agglomerated at the genus level and converted into a presence–absence matrix, where each genus was considered present if detected in a sample and absent otherwise. Inflammation status variables (fibrinogen and haptoglobin) were coded as binary factors, with the normal group used as the reference level.

For each genus, bias-reduced logistic regression was performed using the brglm2 package to estimate the odds of genus presence in animals with elevated fibrinogen or haptoglobin status. Only genera detected in at least five samples and exhibiting variation in presence–absence were retained for analysis to reduce instability associated with sparse taxa. Unadjusted models were fitted with inflammation status as the sole predictor. Bias-reduced estimation was employed to mitigate small-sample bias and address potential separation arising from sparse microbiome data. Odds ratios (ORs) were obtained by exponentiating model coefficients, and 95% confidence intervals (CIs) were calculated using the Wald approximation based on standard errors. *P*-values were derived from model summaries and corrected for multiple testing using the Benjamini–Hochberg false discovery rate (FDR) correction.

To determine whether inflammation-associated keystone taxa identified in the present study have been previously linked to inflammatory processes across different host systems, a literature-based annotation was performed. For each genus, relevant published literature was curated through PubMed searches using the query format *(specified taxon) AND inflammation*, applying the Boolean operator *AND*, to identify reported associations with host inflammation, immune modulation, or functional roles related to gut health and metabolism. Evidence was compiled across cattle, human, and rodent systems and encoded in a binary presence–absence format. Taxa were further categorized based on reported roles (e.g., anti-inflammatory, pro-inflammatory, context dependent- where function varies across host model and unknown- where supporting literature was lacking). These curated annotations were assembled into a reference table and visualized as a heatmap, enabling comparative assessment of keystone taxonomic distribution and inflammation-associated functional patterns across host systems and inflammation statuses.

## DATA AVAILABILITY

All supporting supplementary tables, datafiles, and figures described in this manuscript have been made available via a publicly accessible repository, DOI: 10.5281/zenodo.19546916 (Das et al., 2026). Raw sequencing data and sample metadata can be accessed via the National Center for Biotechnology Information (NCBI) Sequence Read Archive (SRA) under BioProject PRJNA1359400. Codes generated in this analysis are available at github-https://github.com/SlizovskiyLab/Cow_Inflammation_Study.

## ETHICAL CONDUCT OF RESEARCH

All experimental procedures involving animal use were reviewed and approved by the Purdue University Animal Care and Use Committee (Protocol # 0123002347).

## ACKNOWLEDGEMENTS

Financial support for this work was provided by the AnalytixIN faculty fellowship program and Purdue University.

## AUTHOR CONTRIBUTIONS

**Conceptualization:** R.C.N., I.B.S.; **Methodology**: N.T., R.C.N., L.D., D.G.P.N, I.B.S.; **Implementation of computer code and supporting algorithms**: L.D., D.G.P.N., A.D.K., I.B.S.; **Validation**: D.G.P.N., I.B.S.; **Formal Analysis**: L.D., D.G.P.N., A.D.K., I.B.S.; **Study conduct**: N.T., R.C.N, L.D., D.G.P.N., I.B.S.; **Resources**: R.C.N., I.B.S.; **Data curation**: L.D., D.G.P.N., A.D.K.; **Original draft preparation**: L.D., D.G.P.N., I.B.S.; **Reviewing and editing**: L.D., D.G.P.N., N.T., A.D.K., R.C.N., I.B.S.; **Visualizations**: L.D., D.G.P.N., I.B.S.; **Supervision**: R.C.N., I.B.S.; **Project administration**: L.D., R.C.N., I.B.S.; **Funding acquisition**: R.C.N., I.B.S.

## ABBREVIATIONS

DIM1: Day in Milk 1
DIM3: Day in Milk 3
DIM: Day in Milk
DNA: Deoxyribonucleic Acid
16S rRNA: 16S Ribosomal Ribonucleic Acid
rRNA: Ribosomal Ribonucleic Acid
RP: Retained Placenta
IL-6: Interleukin-6
IL-10: Interleukin-10
IFN-γ: Interferon-gamma
IL-1β: Interleukin-1 beta
APPs: Acute Phase Proteins
Hp: Haptoglobin
Fb: Fibrinogen
AGP: Alpha-1-Acid Glycoprotein
ITIH4: Inter-alpha-Trypsin Inhibitor Heavy Chain 4
CRP: C-Reactive Protein
SAA: Serum Amyloid A
LBP: Lipopolysaccharide-Binding Protein
LPS: Lipopolysaccharides
pH: Potential (Power) of Hydrogen
PBS: Phosphate-Buffered Saline
Gb: Gigabase(s)
ASV: Amplicon Sequence Variant
qPCR: Quantitative Polymerase Chain Reaction
FDR: False Discovery Rate
ANOVA: Analysis of Variance
R²: Coefficient of Determination
NMDS: Non-metric Multidimensional Scaling
DADA2: Divisive Amplicon Denoising Algorithm 2
SILVA: SILVA Ribosomal RNA Reference Database
gDNA: Genomic DNA
PCoA: Principal Coordinates Analysis
PERMANOVA: Permutational Multivariate Analysis of Variance
BCS: Body Condition Score
log₂FC: Log2 Fold Change
LOESS: Locally Estimated Scatterplot Smoothing
FE: Fibrinogen Elevated
FN: Fibrinogen Normal
HE: Haptoglobin Elevated
HN: Haptoglobin Normal
BE: Both Elevated (Fibrinogen and Haptoglobin)
BN: Both Normal (Fibrinogen and Haptoglobin)
SpiecEasi: Sparse InversE Covariance Estimation for Ecological Association Inference
SNPI: Standardized Neighborhood Participation Index
OR: Odds Ratio
CI: Confidence Interval
NCBI: National Center for Biotechnology Information
MB: Megabyte(s)
Cat No.: Catalog Number
UCG: Uncultured Group
α: Alpha (significance threshold)
ρ (Rho): Spearman Rank Correlation Coefficient

## LIST OF SUPPLEMENTARY FIGURES AND CAPTIONS

**Supplementary Figure 1: Schematic overview of sample processing and analytical workflow.** Fecal and blood samples were collected from dairy cows during the early postpartum period (DIM1, 3 and 7); DNA was extracted from feces samples and subjected to 16S rRNA gene amplicon sequencing. Raw sequencing reads were processed using the DADA2 pipeline to perform quality filtering, denoising, chimera removal, and amplicon sequence variant (ASV) inference. Taxa assigned by comparing with Silva database. Potential contaminants were identified and removed using *decontam*, informed by negative (extraction, reagent, environmental) controls. SourceTracker was used to estimate the contribution of negative controls sources to fecal microbial communities. Downstream analyses included diversity analyses, differential taxa identification (DeSeq2), microbial abundance network analysis (SPIEC-EASI).

**Supplementary Figure 2: Identification and distribution of contaminant taxa across sample types.** Genus-level contaminant ASVs were identified and quantified using two complementary approaches. (A) *Decontam* was applied to sequencing data to detect putative contaminants based on their prevalence and/or frequency across negative controls. (B) SourceTracker was used to estimate the proportional contribution of potential contamination sources to each sample. The abundance of contaminant genera was calculated and visualized across fecal samples, MockFecal, MockLog.

**Supplementary Figure 3: Module-weighted keystone taxa across inflammation states.** Heatmap showing keystone taxa identified in microbial microbial co-occurance network. Rows represent individual taxa, and columns represent inflammation states: FN = Fibrinogen Normal, HN = Haptoglobin Normal, BN = Both Normal, FE = Fibrinogen Elevated, HE = Haptoglobin Elevated, BE = Both Elevated. Color intensity indicates the number of network modules in which a taxon was identified as a keystone within each condition: white = not a keystone, light blue = single-module keystone, medium blue = keystone in 2 modules, dark blue = keystone in ≥3 modules.

## LIST OF SUPPLEMENTARY TABLES AND CAPTIONS

**Supplementary Table S1: Read statistics for all samples following DADA2 processing.** Summaries of sequencing read counts for each sample at successive stages of the DADA2 pipeline, including input reads, filtered reads, denoised forward and reverse reads, merged reads, and non-chimeric reads. The final read retention (%) represents the proportion of subsampled raw reads retained after chimera removal.

**Supplementary Table S2**: Linear mixed-effects model results for alpha-diversity metrics (Observed richness, Shannon diversity, Simpson index, and Evenness) by fibrinogen, haptoglobin, and combined inflammation status.

**Supplementary Table S3: ASV level differentially abundant taxa associated with inflammation status**. Differentially abundant ASVs in cows with normal versus elevated inflammation, measured by Haptoglobin, Fibrinogen, and Both markers. Only ASVs with significant differential abundance (|log₂FC| ≥ 2, *P*_adj_ < 0.01) are included and grouped into genera. For each genus, the table shows its taxonomic class and phylum, the number of ASVs assigned (n_ASVs), and the number of keystone ASVs (n_Keystone) identified from co-occurrence network analysis. The Keystone column indicates whether any ASV in the genus is a keystone taxon (TRUE/FALSE).

**Supplementary Table S4**: **Details of Keystone ASVs and taxonomic assignments across network modules**. Keystone ASVs identified from co-occurrence network analysis in cows with normal and elevated inflammation as marked by inflammatory markers (Hp, Fb and both). For each ASV, the table reports network topology metrics (degree and eigenvector centrality), taxonomic classification (phylum to species, where available), and module assignment. Only ASVs classified as keystone taxa within their respective networks are shown.

**Supplementary Table S5**: **Taxa showing Occurrence-based odds ratio analysis of keystone microbial genera associated with inflammation status**. Odds ratios (ORs), 95% confidence intervals (CIs), *P*-values, and false discovery rate (FDR)-adjusted *P*-values are shown for non-keystone genera identified through bias-reduced logistic regression analysis. Models evaluated the association between genus presence–absence and elevated inflammation status. Genera with confidence intervals not crossing OR = 1 were considered to exhibit notable directional associations with inflammation status. Odds ratios greater than 1 indicate increased odds of occurrence in elevated inflammation samples, whereas odds ratios less than 1 indicate reduced odds of occurrence.

**Supplementary Table S6**: **Taxa showing occurrence-based odds ratio analysis of non-keystone microbial genera associated with inflammation status**. Odds ratios (OR), 95% confidence intervals (CI), raw *P*-values, and false discovery rate (FDR)-adjusted *P*-values were estimated using presence/absence data for non-keystone genera. OR values > 1 indicate increased odds of occurrence, whereas OR values < 1 indicate decreased odds of occurrence in cows with elevated inflammation status.

## LIST OF SUPPLEMENTARY DATAFILES AND CAPTIONS

**Supplementary datafile 1:** Sample metadata for all sequenced samples. Metadata include sample identifier, sample type, body condition score (BCS), parity, extraction date, plasma fibrinogen (Fb), and haptoglobin (Hp) concentration, inflammation status based on Fb, Hp, and combined inflammation group, 16S rRNA gene copy number, reads at various sequencing steps.

**Supplementary datafile 2:** ASV table for all sequenced samples. Rows represent ASV and columns represent samples. Cell values correspond to the abundance counts of each ASV within each sample. Sample identifiers correspond to those provided in Supplementary Datafile 1.

**Supplementary datafile 3:** Taxonomic classification table for all ASVs detected across sequenced samples. Rows represent ASVs and columns represent sequence, taxonomic ranks, including Kingdom, Phylum, Class, Order, Family, Genus, and Species where available. Taxonomic assignments were generated using the SILVA database (v138.2).

**Supplementary datafile 4**: Mixed-effects model outputs for alpha-diversity analyses. Results of linear mixed-effects models assessing associations between inflammation status and alpha-diversity metrics (Observed richness, Chao1, Shannon diversity, Simpson diversity, and Pielou’s evenness) while accounting for parity group, body condition score (BCS), days in milk (DIM), extraction date, and repeated sampling of individual cows. Model coefficients, standard errors, test statistics, degrees of freedom, raw and FDR-adjusted *P*-values, and marginal and conditional R² values are provided for all models.

**Supplementary datafile 5:** Results of permutational multivariate analysis of variance (PERMANOVA) assessing associations between inflammation status and microbial beta-diversity metrics (Bray-Curtis, weighted UniFrac, unweighted UniFrac, and Aitchison distances) while accounting for parity group, body condition score (BCS), days in milk (DIM), and extraction date. Degrees of freedom, sums of squares, proportion of variance explained (R²), raw and FDR-adjusted *P*-values are provided for all model terms across distance metrics.

**Supplementary datafile 6:** Results of ASV-level differential abundance analysis for haptoglobin-defined inflammation status. For each ASV, normalized mean abundance (baseMean), log2 fold change, standard error of the log2 fold change (lfcSE), test statistic, raw *P*-value, FDR-adjusted *P*-value, nucleotide sequence, and taxonomic classification are provided. Positive log2 fold changes indicate higher abundance in cows with elevated haptoglobin status, whereas negative values indicate lower abundance relative to cows with normal haptoglobin status.

**Supplementary datafile 7:** Results of ASV-level differential abundance analysis for fibrinogen-defined inflammation status. For each ASV, normalized mean abundance (baseMean), log2 fold change, standard error of the log2 fold change (lfcSE), test statistic, raw *P*-value, FDR-adjusted *P*-value, nucleotide sequence, and taxonomic classification are provided. Positive log2 fold changes indicate higher abundance in cows with elevated haptoglobin status, whereas negative values indicate lower abundance relative to cows with normal haptoglobin status.

**Supplementary datafile 8:** Results of ASV-level differential abundance analysis for combined inflammation status. For each ASV, normalized mean abundance (baseMean), log2 fold change, standard error of the log2 fold change (lfcSE), test statistic, raw *P*-value, FDR-adjusted *P*-value, nucleotide sequence, and taxonomic classification are provided. Positive log2 fold changes indicate higher abundance in cows with elevated haptoglobin status, whereas negative values indicate lower abundance relative to cows with normal haptoglobin status.

**Supplementary datafile 9:** Results of ASV-level Spearman correlation analysis between microbial taxa and serum haptoglobin concentrations. For each ASV, Spearman’s correlation coefficient (ρ), raw and FDR-adjusted correlation *P*-values, differential abundance statistics (log2 fold change, standard error, test statistic, baseMean abundance, and adjusted *P*-value), nucleotide sequence, and taxonomic classification are provided. Positive correlation coefficients indicate ASVs positively associated with haptoglobin concentrations, whereas negative coefficients indicate inverse associations.

**Supplementary datafile 10:** Results of ASV-level Spearman correlation analysis between microbial taxa and serum fibrinogen concentrations. For each ASV, Spearman’s correlation coefficient (ρ), raw and FDR-adjusted correlation *P*-values, differential abundance statistics (log2 fold change, standard error, test statistic, baseMean abundance, and adjusted *P*-value), nucleotide sequence, and taxonomic classification are provided. Positive correlation coefficients indicate ASVs positively associated with haptoglobin concentrations, whereas negative coefficients indicate inverse associations.

**Supplementary datafile 11:** Results of ASV-level Spearman correlation analyses between microbial taxa and serum haptoglobin and fibrinogen concentrations. For each amplicon sequence variant (ASV), Spearman’s correlation coefficients (ρ) and corresponding *P*-values for both biomarkers are provided, together with differential abundance statistics, including normalized mean abundance (baseMean), log2 fold change, standard error, test statistic, raw *P*-value, and FDR-adjusted *P*-value. Nucleotide sequences and taxonomic classifications are also included. Positive correlation coefficients indicate ASVs positively associated with biomarker concentrations, whereas negative coefficients indicate inverse associations.

**Supplementary datafile 12:** Candidate keystone ASVs identified from microbial co-occurrence network analyses stratified by inflammation status. For each ASV, inflammation marker classification (haptoglobin or fibrinogen), inflammation status group, taxonomic assignment, network degree, eigenvector centrality, log-transformed degree and eigenvector centrality values, and module membership are provided. Taxa exhibiting extreme values (top 2.5%) for both degree and eigenvector centrality were classified as candidate keystone ASVs.

## REFERENCES

Agudelo, J., & Miller, A. W. (n.d.). Impact of study design, contamination, and data characteristics on results and interpretation of microbiome studies. mSystems, 10(9), e00408–25. 10.1128/msystems.00408-25

Andrews, S. (2010). *FastQC: a quality control tool for high throughput sequence data.* http://www.bioinformatics.babraham.ac.uk/projects/fastqc

Antonov, M., Csárdi, G., Horvát, S., Müller, K., Nepusz, T., Noom, D., Salmon, M., Traag, V., Welles, B. F., & Zanini, F. (2023). *Igraph enables fast and robust network analysis across programming languages* (Version 1). arXiv. 10.48550/ARXIV.2311.10260

Arnalot, L., Pascal, G., Cauquil, L., Vanbergue, E., Foucras, G., & Zened, A. (2025). The bacterial faecal microbiota shifts during the transition period in dairy cows. Animal Microbiome, 7(1), 79. 10.1186/s42523-025-00443-7

Baumgard, L. H., & Rhoads, R. P. (2013). Effects of Heat Stress on Postabsorptive Metabolism and Energetics. Annual Review of Animal Biosciences, 1(1), 311–337. 10.1146/annurev-animal-031412-103644

Bionaz, M., Trevisi, E., Calamari, L., Librandi, F., Ferrari, A., & Bertoni, G. (2007). Plasma Paraoxonase, Health, Inflammatory Conditions, and Liver Function in Transition Dairy Cows. Journal of Dairy Science, 90(4), 1740–1750. 10.3168/jds.2006-445

Blondel, V. D., Guillaume, J.-L., Lambiotte, R., & Lefebvre, E. (2008). Fast unfolding of communities in large networks. Journal of Statistical Mechanics: Theory and Experiment, 2008(10), P10008. 10.1088/1742-5468/2008/10/P10008

Bolger, A. M., Lohse, M., & Usadel, B. (2014). Trimmomatic: A flexible trimmer for Illumina sequence data. Bioinformatics, 30(15), 2114–2120. 10.1093/bioinformatics/btu170

Bradford, B. J., Yuan, K., Farney, J. K., Mamedova, L. K., & Carpenter, A. J. (2015a). Invited review: Inflammation during the transition to lactation: New adventures with an old flame. Journal of Dairy Science, 98(10), 6631–6650. 10.3168/jds.2015-9683

Bradford, B. J., Yuan, K., Farney, J. K., Mamedova, L. K., & Carpenter, A. J. (2015b). *Invited review:* Inflammation during the transition to lactation: New adventures with an old flame. Journal of Dairy Science, 98(10), 6631–6650. 10.3168/jds.2015-9683

Brown, W. E., & Bradford, B. J. (2021). Invited review: Mechanisms of hypophagia during disease. Journal of Dairy Science, 104(9), 9418–9436. 10.3168/jds.2021-20217

Bruinjé, T. C., & LeBlanc, S. J. (2025). Invited Review: Inflammation and Health in the Transition Period Influence Reproductive Function in Dairy Cows. Animals, 15(5), 633. 10.3390/ani15050633

Callahan, B. J., McMurdie, P. J., Rosen, M. J., Han, A. W., Johnson, A. J. A., & Holmes, S. P. (2016). DADA2: High-resolution sample inference from Illumina amplicon data. Nature Methods, 13(7), 581–583. 10.1038/nmeth.3869

Castillo-Lopez, E., Ricci, S., Rivera-Chacon, R., Sener-Aydemir, A., Pacífico, C., Reisinger, N., Schwartz-Zimmermann, H. E., Berthiller, F., Kreuzer-Redmer, S., & Zebeli, Q. (2024). Dynamic interplay of immune response, metabolome, and microbiota in cows during high-grain feeding: Insights from multi-omics analysis. Microbiology Spectrum, 12(10), e00944–24. 10.1128/spectrum.00944-24

Clemente, J. C., Manasson, J., & Scher, J. U. (2018). The role of the gut microbiome in systemic inflammatory disease. *BMJ*, j5145. 10.1136/bmj.j5145

Das, L., Puerres Narvaez, D. G., Taechachokevivat, N., Kimball, A. D., Neves, R., & Slizovskiy, I. (2026). *Dataset and Supplementary Materials for: Postpartum Systemic Inflammation is Reflected in Early Distinct Fecal Microbiome Differences in Dairy Cows* [Dataset]. Zenodo. 10.5281/ZENODO.19546916

Davis, N. M., Proctor, D. M., Holmes, S. P., Relman, D. A., & Callahan, B. J. (2018). Simple statistical identification and removal of contaminant sequences in marker-gene and metagenomics data. Microbiome, 6(1), 226. 10.1186/s40168-018-0605-2

De Filippo, C., Cavalieri, D., Di Paola, M., Ramazzotti, M., Poullet, J. B., Massart, S., Collini, S., Pieraccini, G., & Lionetti, P. (2010). Impact of diet in shaping gut microbiota revealed by a comparative study in children from Europe and rural Africa. Proceedings of the National Academy of Sciences, 107(33), 14691–14696. 10.1073/pnas.1005963107

Dean, C. J., Deng, Y., Wehri, T. C., Pena-Mosca, F., Ray, T., Crooker, B. A., Godden, S. M., Caixeta, L. S., & Noyes, N. R. (2024). The impact of kit, environment, and sampling contamination on the observed microbiome of bovine milk. mSystems, 9(6), e01158–23. 10.1128/msystems.01158-23

Díaz, S., Escobar, J. S., & Avila, F. W. (2021). Identification and Removal of Potential Contaminants in 16S rRNA Gene Sequence Data Sets from Low-Microbial-Biomass Samples: An Example from Mosquito Tissues. mSphere, 6(3), e00506–21. 10.1128/mSphere.00506-21

Dunière, L., Esparteiro, D., Lebbaoui, Y., Ruiz, P., Bernard, M., Thomas, A., Durand, D., Forano, E., & Chaucheyras-Durand, F. (2021). Changes in Digestive Microbiota, Rumen Fermentations and Oxidative Stress around Parturition Are Alleviated by Live Yeast Feed Supplementation to Gestating Ewes. Journal of Fungi, 7(6), 447. 10.3390/jof7060447

Forman-Ankjær, B., Hvid-Jensen, F., Kobel, C. M., & Greve, T. (2024). Short communication: First case of bacteraemia caused by *Dielma fastidiosa* in a patient hospitalized with diverticulitis. APMIS, 132(2), 130–133. 10.1111/apm.13367

Gohl, D. M., Vangay, P., Garbe, J., MacLean, A., Hauge, A., Becker, A., Gould, T. J., Clayton, J. B., Johnson, T. J., Hunter, R., Knights, D., & Beckman, K. B. (2016). Systematic improvement of amplicon marker gene methods for increased accuracy in microbiome studies. Nature Biotechnology, 34(9), 942–949. 10.1038/nbt.3601

Gragg, S. E., Loneragan, G. H., Nightingale, K. K., Brichta-Harhay, D. M., Ruiz, H., Elder, J. R., Garcia, L. G., Miller, M. F., Echeverry, A., Ramírez Porras, R. G., & Brashears, M. M. (2013). Substantial within-Animal Diversity of Salmonella Isolates from Lymph Nodes, Feces, and Hides of Cattle at Slaughter. Applied and Environmental Microbiology, 79(15), 4744–4750. 10.1128/AEM.01020-13

Guimerà, R., & Nunes Amaral, L. A. (2005). Functional cartography of complex metabolic networks. Nature, 433(7028), 895–900. 10.1038/nature03288

Guo, C., Liu, J., Wei, Y., Du, W., & Li, S. (2024). Comparison of the gastrointestinal bacterial microbiota between dairy cows with and without mastitis. Frontiers in Microbiology, 15, 1332497. 10.3389/fmicb.2024.1332497

He, B., Nohara, K., Ajami, N. J., Michalek, R. D., Tian, X., Wong, M., Losee-Olson, S. H., Petrosino, J. F., Yoo, S.-H., Shimomura, K., & Chen, Z. (2015). Transmissible microbial and metabolomic remodeling by soluble dietary fiber improves metabolic homeostasis. Scientific Reports, 5(1), 10604. 10.1038/srep10604

Honda, K., & Littman, D. R. (2012). The Microbiome in Infectious Disease and Inflammation. Annual Review of Immunology, 30(1), 759–795. 10.1146/annurev-immunol-020711-074937

Huang, S., Ji, S., Wang, F., Huang, J., Alugongo, G. M., & Li, S. (2020). Dynamic changes of the fecal bacterial community in dairy cows during early lactation. AMB Express, 10(1), 167. 10.1186/s13568-020-01106-3

Huzzey, J. M., Nydam, D. V., Grant, R. J., & Overton, T. R. (2011). Associations of prepartum plasma cortisol, haptoglobin, fecal cortisol metabolites, and nonesterified fatty acids with postpartum health status in Holstein dairy cows. Journal of Dairy Science, 94(12), 5878–5889. 10.3168/jds.2010-3391

International Dairy Federation. (2025). *IDF Annual Report 2024–*2025 [Annual Report]. https://zuivel-nl.files.svdcdn.com/production/images/IDF-Annual-Report-2024-2025.pdf

Ishikawa, Y., Nakada, K., Hagiwara, K., Kirisawa, R., Iwai, H., Moriyoshi, M., & Sawamukai, Y. (2004). Changes in Interleukin-6 Concentration in Peripheral Blood of Pre- and Post-Partum Dairy Cattle and Its Relationship to Postpartum Reproductive Diseases. Journal of Veterinary Medical Science, 66(11), 1403–1408. 10.1292/jvms.66.1403

Jahan, N., Trevisi, E., Minuti, A., Ahmed, S., & Haque, M. N. (2025). Pro-inflammatory cytokines as indicators of health status around calving in dairy cows. SSRN. 10.2139/ssrn.5938823

Jia, Y., Zhao, S., Guo, W., Peng, L., Zhao, F., Wang, L., Fan, G., Zhu, Y., Xu, D., Liu, G., Wang, R., Fang, X., Zhang, H., Kristiansen, K., Zhang, W., & Chen, J. (2022). Sequencing introduced false positive rare taxa lead to biased microbial community diversity, assembly, and interaction interpretation in amplicon studies. Environmental Microbiome, 17(1), 43. 10.1186/s40793-022-00436-y

Jiang, P., Yu, F., Zhou, X., Shi, H., He, Q., & Song, X. (2024). Dissecting causal links between gut microbiota, inflammatory cytokines, and DLBCL: A Mendelian randomization study. Blood Advances, 8(9), 2268–2278. 10.1182/bloodadvances.2023012246

Johnzon, C.-F., Dahlberg, J., Gustafson, A.-M., Waern, I., Moazzami, A. A., Östensson, K., & Pejler, G. (2018). The Effect of Lipopolysaccharide-Induced Experimental Bovine Mastitis on Clinical Parameters, Inflammatory Markers, and the Metabolome: A Kinetic Approach. Frontiers in Immunology, 9, 1487. 10.3389/fimmu.2018.01487

Kagambèga, A., Lienemann, T., Aulu, L., Traoré, A. S., Barro, N., Siitonen, A., & Haukka, K. (2013). Prevalence and characterization of Salmonella enterica from the feces of cattle, poultry, swine and hedgehogs in Burkina Faso and their comparison to human Salmonella isolates. BMC Microbiology, 13(1), 253. 10.1186/1471-2180-13-253

Kerwin, A. L., Burhans, W. S., Mann, S., Nydam, D. V., Wall, S. K., Schoenberg, K. M., Perfield, K. L., & Overton, T. R. (2022). Transition cow nutrition and management strategies of dairy herds in the northeastern United States: Part II—Associations of metabolic- and inflammation-related analytes with health, milk yield, and reproduction. Journal of Dairy Science, 105(6), 5349–5369. 10.3168/jds.2021-20863

Khafipour, E., Li, S., Plaizier, J. C., & Krause, D. O. (2009). Rumen Microbiome Composition Determined Using Two Nutritional Models of Subacute Ruminal Acidosis. Applied and Environmental Microbiology, 75(22), 7115–7124. 10.1128/AEM.00739-09

Knights, D., Kuczynski, J., Charlson, E. S., Zaneveld, J., Mozer, M. C., Collman, R. G., Bushman, F. D., Knight, R., & Kelley, S. T. (2011). Bayesian community-wide culture-independent microbial source tracking. Nature Methods, 8(9), 761–763. 10.1038/nmeth.1650

Kodera, S. M., Das, P., Gilbert, J. A., & Lutz, H. L. (2022). Conceptual strategies for characterizing interactions in microbial communities. iScience, 25(2), 103775. 10.1016/j.isci.2022.103775

Ku, J.-Y., Lee, M.-J., Jung, Y., Choi, H.-J., & Park, J. (2025). Changes in the gut microbiome due to diarrhea in neonatal Korean indigenous calves. Frontiers in Microbiology, 16, 1511430. 10.3389/fmicb.2025.1511430

Kumar, M., Babaei, P., Ji, B., & Nielsen, J. (2016). Human gut microbiota and healthy aging: Recent developments and future prospective. Nutrition and Healthy Aging, 4(1), 3–16. 10.3233/NHA-150002

Kurtz, Z. D., Müller, C. L., Miraldi, E. R., Littman, D. R., Blaser, M. J., & Bonneau, R. A. (2015). Sparse and Compositionally Robust Inference of Microbial Ecological Networks. PLOS Computational Biology, 11(5), e1004226. 10.1371/journal.pcbi.1004226

Kvidera, S. K., Horst, E. A., Abuajamieh, M., Mayorga, E. J., Fernandez, M. V. S., & Baumgard, L. H. (2017). Glucose requirements of an activated immune system in lactating Holstein cows. Journal of Dairy Science, 100(3), 2360–2374. 10.3168/jds.2016-12001

Leo Lahti [Aut, C. (2017). *Microbiome* [Computer software]. Bioconductor. 10.18129/B9.BIOC.MICROBIOME

Li, C., Stražar, M., Mohamed, A. M. T., Pacheco, J. A., Walker, R. L., Lebar, T., Zhao, S., Lockart, J., Dame, A., Thurimella, K., Jeanfavre, S., Brown, E. M., Ang, Q. Y., Berdy, B., Sergio, D., Invernizzi, R., Tinoco, A., Pishchany, G., Vasan, R. S., … Xavier, R. J. (2024). Gut microbiome and metabolome profiling in Framingham heart study reveals cholesterol-metabolizing bacteria. Cell, 187(8), 1834–1852.e19. 10.1016/j.cell.2024.03.014

Li, F., Wang, M., Wang, J., Li, R., & Zhang, Y. (2019). Alterations to the Gut Microbiota and Their Correlation With Inflammatory Factors in Chronic Kidney Disease. Frontiers in Cellular and Infection Microbiology, 9, 206. 10.3389/fcimb.2019.00206

Liu, J., Ahmad, A. A., Yang, C., Zhang, J., Zheng, J., Liang, Z., Wang, F., Zhai, H., Qin, S., Yang, F., & Ding, X. (2024). Modulations in gastrointestinal microbiota during postpartum period fulfill energy requirements and maintain health of lactating Tibetan cattle. Frontiers in Microbiology, 15, 1369173. 10.3389/fmicb.2024.1369173

Love, M. I., Huber, W., & Anders, S. (2014). Moderated estimation of fold change and dispersion for RNA-seq data with DESeq2. Genome Biology, 15(12), 550. 10.1186/s13059-014-0550-8

Lun, H., Yang, W., Zhao, S., Jiang, M., Xu, M., Liu, F., & Wang, Y. (2019). Altered gut microbiota and microbial biomarkers associated with chronic kidney disease. MicrobiologyOpen, 8(4), e00678. 10.1002/mbo3.678

McLaren, M. R., Willis, A. D., & Callahan, B. J. (2019). Consistent and correctable bias in metagenomic sequencing experiments. eLife, 8, e46923. 10.7554/eLife.46923

McMurdie, P. J., & Holmes, S. (2013). phyloseq: An R Package for Reproducible Interactive Analysis and Graphics of Microbiome Census Data. PLoS ONE, 8(4), e61217. 10.1371/journal.pone.0061217

Mekuriaw, Y. (2023). Negative energy balance and its implication on productive and reproductive performance of early lactating dairy cows: Review paper. Journal of Applied Animal Research, 51(1), 220–228. 10.1080/09712119.2023.2176859

Millar, H. R., Simpson, J. G., & Stalker, A. L. (1971). An evaluation of the heat precipitation method for plasma fibrinogen estimation. Journal of Clinical Pathology, 24(9), 827–830. 10.1136/jcp.24.9.827

Murray, R. D., Downham, D. Y., Demirkan, I., & Carter, S. D. (2002). Some relationships between spirochaete infections and digital dermatitis in four UK dairy herds. Research in Veterinary Science, 73(3), 223–230. 10.1016/S0034-5288(02)00027-9

Newman, M. E. J. (2003). Mixing patterns in networks. Physical Review E, 67(2), 026126. 10.1103/PhysRevE.67.026126

Ni, J., Wu, G. D., Albenberg, L., & Tomov, V. T. (2017). Gut microbiota and IBD: Causation or correlation? Nature Reviews Gastroenterology & Hepatology, 14(10), 573–584. 10.1038/nrgastro.2017.88

Oksanen, J., Simpson, G. L., Blanchet, F. G., Kindt, R., Legendre, P., Minchin, P. R., O’Hara, R. B., Solymos, P., Stevens, M. H. H., Szoecs, E., Wagner, H., Barbour, M., Bedward, M., Bolker, B., Borcard, D., Borman, T., Carvalho, G., Chirico, M., De Caceres, M., … Weedon, J. (2001). vegan: Community Ecology Package (p. 2.7–2) [Dataset]. 10.32614/CRAN.package.vegan

Paulina, J. & Stefaniak Tadeusz. (2011). Acute Phase Proteins in Cattle. Unpublished. 10.13140/2.1.4279.9042

Plaizier, J. C., Khafipour, E., Li, S., Gozho, G. N., & Krause, D. O. (2012). Subacute ruminal acidosis (SARA), endotoxins and health consequences. Animal Feed Science and Technology, 172(1–2), 9–21. 10.1016/j.anifeedsci.2011.12.004

Quast, C., Pruesse, E., Yilmaz, P., Gerken, J., Schweer, T., Yarza, P., Peplies, J., & Glöckner, F. O. (2012). The SILVA ribosomal RNA gene database project: Improved data processing and web-based tools. Nucleic Acids Research, 41(D1), D590–D596. 10.1093/nar/gks1219

Reczyńska, D., Zalewska, M., Czopowicz, M., Kaba, J., Zwierzchowski, L., & Bagnicka, E. (2018). Acute Phase Protein Levels as An Auxiliary Tool in Diagnosing Viral Diseases in Ruminants—A Review. Viruses, 10(9), 502. 10.3390/v10090502

Saco, Y., & Bassols, A. (2023). Acute phase proteins in cattle and swine: A review. Veterinary Clinical Pathology, 52(S1), 50–63. 10.1111/vcp.13220

Saco, Y., Crusellas-Villorbina, N., Peña, R., Pato, R., Marti, S., Pisoni, L., Devant, M., Pelegrí-Pineda, A., & Bassols, A. (2025). Bovine fecal biomarkers of intestinal inflammatory process: Calprotectin and lactoferrin, a comparative study. Research in Veterinary Science, 183, 105500. 10.1016/j.rvsc.2024.105500

Saleem, F., Ametaj, B. N., Bouatra, S., Mandal, R., Zebeli, Q., Dunn, S. M., & Wishart, D. S. (2012). A metabolomics approach to uncover the effects of grain diets on rumen health in dairy cows. Journal of Dairy Science, 95(11), 6606–6623. 10.3168/jds.2012-5403

Serbetci, I., González-Grajales, L. A., Herrera, C., Ibanescu, I., Tekin, M., Melean, M., Magata, F., Malama, E., Bollwein, H., & Scarlet, D. (2024). Impact of negative energy balance and postpartum diseases during the transition period on oocyte quality and embryonic development in dairy cows. Frontiers in Veterinary Science, 10. 10.3389/fvets.2023.1328700

Sheldon, I. M., Cronin, J., Goetze, L., Donofrio, G., & Schuberth, H.-J. (2009). Defining Postpartum Uterine Disease and the Mechanisms of Infection and Immunity in the Female Reproductive Tract in Cattle1. Biology of Reproduction, 81(6), 1025–1032. 10.1095/biolreprod.109.077370

Stipetic, K., Chang, Y., Peters, K., Salem, A., Doiphode, S. H., McDonough, P. L., Chang, Y. F., Sultan, A., & Mohammed, H. O. (2016). The risk of carriage of *Salmonella* spp. And Listeria monocytogenes in food animals in dynamic populations. Veterinary Medicine and Science, 2(4), 246–254. 10.1002/vms3.39

Trevisi, E., Jahan, N., Bertoni, G., Ferrari, A., & Minuti, A. (2015). Pro-Inflammatory Cytokine Profile in Dairy Cows: Consequences for New Lactation. Italian Journal of Animal Science, 14(3), 3862. 10.4081/ijas.2015.3862

U.S. Department of Agriculture. (2025). https://ers.usda.gov/sites/default/files/_laserfiche/outlooks/37809/56729_oce-2016-1.pdf?v=90927

Wang, L., Wu, D., Zhang, Y., Li, K., Wang, M., & Ma, J. (2023). Dynamic distribution of gut microbiota in cattle at different breeds and health states. Frontiers in Microbiology, 14, 1113730. 10.3389/fmicb.2023.1113730

Wang, S., Kong, F., Dai, D., Li, C., Hao, Y., Wang, E., Cao, Z., Wang, Y., Wang, W., & Li, S. (2025). Deterministic succession patterns in the rumen and fecal microbiome associate with host metabolic shifts in peripartum dairy cattle. GigaScience, 14, giaf042. 10.1093/gigascience/giaf042

Westhoff, T. A., Sipka, A. S., Overton, T. R., Klaessig, S., & Mann, S. (2025). Evaluation of circulating cytokine concentrations and ex vivo indicators of the inflammatory response in transition dairy cows fed pre- and postpartum diets differing in metabolizable protein supply. Journal of Dairy Science, 108(6), 6427–6438. 10.3168/jds.2024-26074

Wickham, H. (2016). Ggplot2. Springer International Publishing. 10.1007/978-3-319-24277-4

Zhang, B., Wang, N., He, G., Chen, T., Zong, J., Shen, C., Wang, Y., Li, C., Yin, X., Meng, Y., Chang, F., Wang, S., Li, C., & Zhou, X. (2025). *AKKERMANSIA MUCINIPHILA* -Derived Outer Membrane Vesicles as a Novel Therapeutic Approach for Mastitis: Insights From In Vitro and Vivo Studies. The FASEB Journal, 39(13), e70770. 10.1096/fj.202500877RR

Zhang, C., Wang, M., Liu, H., Jiang, X., Chen, X., Liu, T., Yin, Q., Wang, Y., Deng, L., Yao, J., & Wu, S. (2023). Multi-omics reveals that the host-microbiome metabolism crosstalk of differential rumen bacterial enterotypes can regulate the milk protein synthesis of dairy cows. Journal of Animal Science and Biotechnology, 14(1), 63. 10.1186/s40104-023-00862-z

Zhao, X., Zhang, Y., Rahman, A., Chen, M., Li, N., Wu, T., Qi, Y., Zheng, N., Zhao, S., & Wang, J. (2024). Rumen microbiota succession throughout the perinatal period and its association with postpartum production traits in dairy cows: A review. Animal Nutrition, 18, 17–26. 10.1016/j.aninu.2024.04.013

Zhong, P., Ren, A., Cui, J., Guo, C., Zhang, Y., Diao, Q., Liu, X., Zhang, N., Tu, Y., & Bi, Y. (2026). Microbial landscapes in dairy cow diseases: From localized dysbiosis to inter-organ axes. Npj Biofilms and Microbiomes. 10.1038/s41522-026-00988-8

